# Cytokinins control secondary cell wall formation in the inflorescence stem of Arabidopsis

**DOI:** 10.1101/2023.07.26.550726

**Authors:** Vojtech Didi, Dominique Arnaud, Anna Pacinková, Radek Jupa, Radim Cegan, Alesia Melnikava, Jana Vasickova, Mariana Benitez, Faride Unda, Tereza Dobisova, Willi Riber, Zuzana Dostalova, Shawn D. Mansfield, Ondrej Novak, Miroslav Strnad, Roman Hobza, Vít Gloser, Eva Budinska, Jan Hejatko

## Abstract

Spatiotemporal control over developmental programs is vital to all organisms. Here we show that cytokinin (signaling) deficiency leads to early secondary cell wall (SCW) formation in Arabidopsis inflorescence stem that associates with precocious upregulation of a SCW transcriptional cascade controlled by NAC TFs (NSTs). We demonstrate that cytokinin signaling through the AHK2/3 and the ARR1/10/12 suppresses the expression of several *NSTs* and SCW formation in the apical portions of stems. Exogenous cytokinin application reconstituted both proper development and apical-basal gradient of *NST1* and *NST3* in a cytokinin biosynthesis-deficient mutant. We show that *AHK2* and *AHK3* required functional *NST1* or *NST3* to control SCW initiation in the interfascicular fibers, further evidencing that cytokinins act upstream of *NST*s transcription factors. The premature onset of a rigid SCW biosynthesis and altered expression of *NST1/3* and *VND6/7* due to cytokinin deficiency led to the formation of smaller tracheary elements (TEs) and impaired hydraulic conductivity. We conclude that cytokinins downregulate *NSTs* to inhibit premature SCW formation in the apical part of the inflorescence stem, facilitating thus the development of fully functional TEs and interfascicular fibers.

**Summary statement:** Cytokinins attenuate premature secondary cell wall (SCW) formation via downregulating the expression of NAC TFs, the master switches of SCW transcriptional cascade, thus affecting the tracheary elements size and conductivity.

## Introduction

In the plant postembryonic development, growth is largely controlled by the activity of apical (shoot and root) and lateral (procambium, cambium and axillary) meristems. Meristems maintain a pool of totipotent stem cells that divide and differentiate, facilitating the formation of new plant organs and tissues (Greb and Lohmann, 2016). Procambium, together with latter differentiating cambium ensure formation of a new vascular tissue that is necessary prerequisite for proper plant growth. In the growing shoot apex, procambium is formed first, followed by phloem and subsequently xylem specification. The newly formed tissue connect with previously formed vasculature, thus ensuring the continuity of the shoot vascular system (Esau, 1977). As a result, the developmental gradient along the apical-basal axis is established (Altamura et al., 2001; Baima et al., 2001).

*Arabidopsis* inflorescence stems show a collateral eustele pattern, where asymmetric cell division of procambial stem cells leads to phloem and xylem specification on the outer and inner side of the procambial layer, respectively. In the xylem, two cell types, protoxylem and metaxylem, are able to form conductive tracheary elements (TEs) embedded in the surrounding xylem parenchyma and xylary fiber cells (Ruzicka et al., 2015; Schuetz et al., 2013). All fully differentiated xylem cells are characterized by the presence of secondary cell walls (SCWs). Older portions (medial and basal internodia) of stems reaching the stage 1, corresponding to first silique formation (Altamura et al., 2001), develop interfascicular fibers (also called interfascicular arcs), characterized by massive secondary wall thickening and spanning the neighboring vascular bundles as a mechanical support.

In contrast to the dynamic structure of primary cell walls, the lignified SCW is rigid, conferring the SCW-possessing cells the resistance to greater mechanical stresses (Zhong and Ye, 2015). This enables xylary and interfascicular fibers to stabilize the plant body and provides TEs with the ability to withstand the negative pressure resulting from the transpirational stream. SCW synthesis permits xylem to fulfill its main functional role of key evolutionary importance, which is water conductance throughout the plant body.

The differentiation of vascular tissues represents a complex process involving dramatic changes in the cell shape and functional properties of the conducting and supporting tissues. Early after specification, the future TEs undergo tremendous expansion in both the radial and longitudinal directions. Once expansion ceases, deposition of the individual components of the SCW follows, culminating in cell death and final maturation of the functional TE (Ruzicka et al., 2015). Xylary and interfascicular fibers undergo a similar differentiation process but remain alive. It is obvious that the entire process requires a delicate spatiotemporal control of several integral processes. In the last few years, significant progress has been made in our understanding of the mechanisms controlling the onset and structure of the SCW (Zhong and Ye, 2015). The transcriptional regulatory cascade was described, allowing the initiation of SCW formation in response to activation by NAM/ATAF/CUC (NAC) transcription factors (Hussey et al., 2013; Taylor-Teeples et al., 2015). NAC transcription factors are considered as master regulators or master switches with VASCULAR-RELATED NAC-DOMAIN 6 (VND6) and VND7 and NAC SECONDARY WALL THICKENING PROMOTING FACTOR 1 (NST1) and NST3 regulating SCW formation in vessels (TEs) and fibers, respectively. When overexpressed, they are able to induce ectopic SCW thickenings even in case of fully differentiated tissues including floral organs or the rosette leaf epidermis (Kubo et al., 2005; Mitsuda et al., 2007). However, how NAC master regulators are controlled in the complex network of developmental events leading to xylem differentiation remains to be clarified.

The plant hormones, cytokinins, are potent regulators of plant development. One of the main roles of cytokinins appears to be the fine control over the equilibrium of cell division and cell differentiation (Dello Ioio et al., 2008). Although cytokinins have long been known to be important for xylogenesis and both procambium and cambium formation (Hejatko et al., 2009; Matsumoto-Kitano et al., 2008; Nieminen et al., 2008; Ye et al., 2021), the specific role(s) of cytokinins in the control of SCW formation are just being elucidated (Didi et al., 2015). Cytokinins are recognized by the sensor ARABIDOPSIS HISTIDINE KINASES (AHKs)2-4 that transmit the signal to the nucleus *via* a multistep phosphorelay (MSP) pathway. The first downstream partners of AHK2, AHK3 and AHK4 are ARABIDOPSIS HIS-CONTAINING PHOSPHOTRANSMITTERS (AHPs)1-6. AHPs transfer the signal to the nucleus, where the final transphosphorylation step occurs with the activation of type-B ARABIDOPISIS RESPONSE REGULATORS (RRBs), the GARP-domain containing transcription factors which facilitate the control over expression of primary cytokinin-regulated genes (Kieber and Schaller, 2018). Among the direct targets of RRBs are the type-A response regulators (*RRAs*), whose expression was previously shown to provide a reliable quantitative measure of cytokinin signaling activity (Gordon et al., 2009; Pernisova et al., 2009).

Herein, we provide evidence that cytokinins control xylem and fiber development in the *Arabidopsis* inflorescence stem, *via* the transcriptional regulation of the genes for NAC transcription factors NST1/NST3 and VND6/VND7, thus ensuring the proper spatiotemporal control of SCW initiation that is critical for the formation of fully functional fibers and TEs.

## Results

### Attenuation of cytokinin signaling and endogenous cytokinin deficiency leads to the premature secondary cell wall formation

In *Arabidopsis* inflorescence stem, the SCW thickening was found to occur after flower anthesis and individual developmental stages were defined as based on the progress of reproductive growth (Altamura et al., 2001). To normalize the differences of developmental stage between individual plants and genotypes, all samples were collected at stage 1, corresponding to the moment of first silique (elongated pistil) formation (Fig. S1A). A clear developmental gradient of SCWs is apparent along the apical/basal axis of inflorescence stem in *Arabidopsis* wild-type (WT) Col-0 plants (Fig. 1). In the apical internode (the youngest internode below the flowers), vascular bundles contain only a few proto- and (rarely) metaxylem cells with a developed SCW, while no lignified (SCW-containing) cells were observed in the interfascicular regions. In contrast, in the basal part of the stem (the oldest internode above the rosette leaves), number of metaxylem cells is fully developed and SCW-containing xylem, xylary and extraxylary fibers are clearly visible in vascular bundles and interfascicular regions, respectively, the latter forming there what is called interfascicular arcs (Fig. 1A). Fully differentiated SCW-containing cells, both in vascular bundles and interfascicular arcs, are characterized by the presence of lignin and xylose, an aromatic polymer and the main component of *Arabidopsis* SCW hemicellulose, respectively (Zhong and Ye, 2015). In agreement with our histological analysis, a gradual increase of acid insoluble lignin and xylose content was apparent from the apical to middle and basal internodes of the inflorescence stem (Fig. 1C).

**Figure 1.**
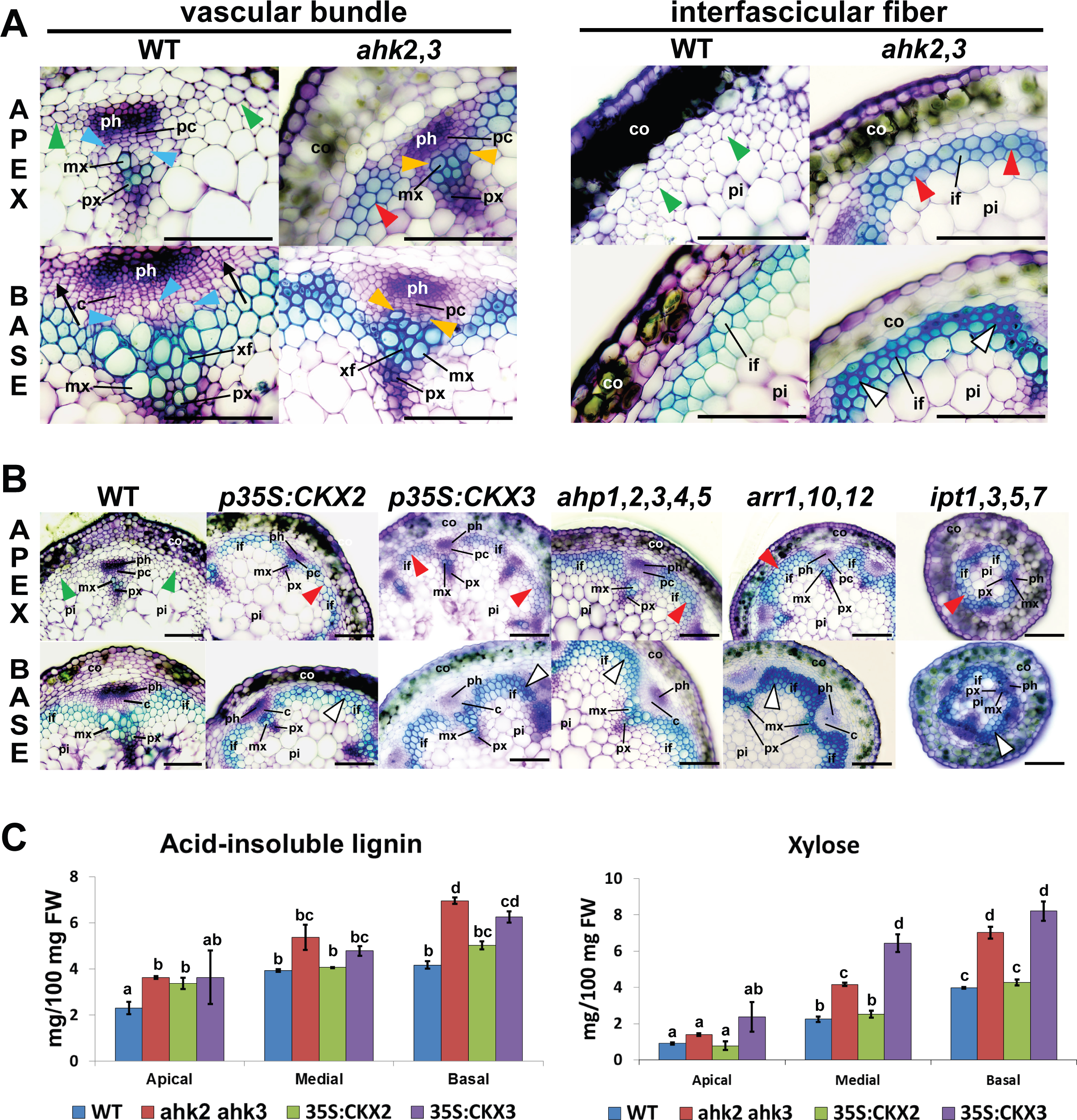
Cytokinin (signaling) deficiency leads to premature SCW formation. **(A)** Toluidine blue-stained transverse section through the living WT Col-0 and *ahk2,3* inflorescence stems showing premature SCW formation (red arrowhead) in the apical internode (APEX) of *ahk2,3*. Note the comparative lack of SCW formation (green arrowhead) in WT plants. Differentiating TEs surrounded by primary CW and SCW are depicted by blue and orange arrowheads, respectively. A stronger toluidine blue staining indicative of increased SCW deposition (empty arrowheads) can also be seen in the basal internode (BASE) of *ahk2,3*, particularly in interfascicular fibers (see also in B). The presence of the interfascicular cambium at the base of the WT (black arrow) indicates the developmental switch to secondary growth that is missing in the *ahk2,3* line. High resolution micrographs are provided allowing to zoom to even 800 % of the original image size to see the detailed VBs structure. c – cambium; co – cortex; if-interfascicular fibers; mx – metaxylem; pc – procambium; ph – phloem; pi – pith; px – protoxylem; xf – xylary fibers. Scale bars 50 µm. **(B)** Premature SCW formation (interfascicular arcs, red arrowhead) in the apex of cytokinin (signaling) deficient plants compared to Col-0 WT (absence of SCW depicted by green arrowhead); abbreviations as in A. Scale bars 50 µm. **(C)** Relative acid-insoluble lignin and xylose content (±SE, three biological replicates) in apical, medial and basal sections of inflorescence stems of WT Col-0, *ahk2,3*, *p35S:CKX2* and *p35S:CKX3*. Data are means ± SE of two independent experiments (n≥3). Different letters indicate significant differences at *P* < 0.05 based on a Tukey’s HSD test.

Cytokinins were identified as one of the key regulators of plant cell differentiation (Dello Ioio et al., 2008). In addition, the role of cytokinins in anther lignification and associated SCW formation has been described (Jung et al., 2008). Among the three cytokinin receptors known to date, AHK2 and AHK3 were shown to be the most important during both the early (Higuchi et al., 2004; Nishimura et al., 2004; Riefler et al., 2006) and later stages of shoot development (Hejatko et al., 2009). When compared with WT (Col-0), we observed a premature onset of SCW formation in the apical portion of inflorescence stems in the *ahk2-1 ahk3-1* [*ahk2,3*; (Nishimura et al., 2004)] double mutant at stage 1 (Fig. 1A). In contrast to WT, where a few TEs (mostly of protoxylem identity) with SCWs were found exclusively in vascular bundles, in *ahk2,3* apices more cells with clearly present SCWs were apparent in both vascular bundles and interfascicular arcs. Further, in WT, the first differentiating TEs that could be distinguished as enlarged cells positioned close to the procambium were still encircled with primary cell wall only (highlighted in Fig. 1A). Compared to that, in both apical and basal internodia of *ahk2,3,* all the cells morphologically distinguishable as differentiating TEs seem to develop SCW, even if located in a very close vicinity to procambium (Fig. 1A), implying premature SCW formation even in the vascular bundles. More intense toluidine blue staining also suggested a greater deposition of SCW materials in the basal internode in *ahk2,3*. Relative to WT, measurements of acid-insoluble lignin and xylose confirmed an increase in the SCW components in all three (apical, middle and basal) internodes of *ahk2,3* (Fig. 1C). Importantly, no ectopic SCW formation (e.g. in the pith or phloem) was observed, suggesting that attenuated cytokinin signaling does not affect the determination of xylem and fiber cell identity.

Similar phenotypes were also observed in other cytokinin signaling deficient lines such as the quintuple mutant *ahp1,2,3,4,5*, the triple *arr1,10,12* and *arr1*,*12* and *arr10*,*12* double mutants (Fig. 1B and Fig. S1B). Furthermore, premature SCW deposition in interfascicular region was also apparent in plants defective in cytokinin biosynthesis (the triple *atipt3,5,7* and particularly the quadruple *atipt1,3,5,7* mutant showing the strongest phenotype), and in plants with depleted endogenous cytokinins due to overexpression of cytokinin catabolizing *CYTOKININ OXIDASE/DEHYDROGENASE* genes (*p35S:CKX2* and *p35S:CKX3*; Fig. 1B, Fig. S1B). Similar to the *ahk2,3*, the overexpression lines, *p35S:CKX2* and *p35S:CKX3*, produced altered acid-insoluble lignin and xylose gradients, mainly due to increased SCW formation in the apical internode (Fig. 1C). The content of other sugars showed differential responses in individual cytokinin (signaling) deficient lines. However, the changes observed in the glucose, galactose, rhamnose, as well as acid-soluble lignin seem to support our data, suggesting the regulatory role of cytokinins in the SCW formation (Fig. S2).

Overall, these data suggest that cytokinins control both xylem and extraxylary cell files (interfascicular arcs) development along the apical-basal axis, possibly acting as negative regulators to prevent precocious SCW formation.

### Cytokinins regulate the transcriptional cascade controlling the onset of secondary cell wall biosynthesis

To elucidate the molecular mechanisms of cytokinin-mediated control over SCW formation, we performed genome-wide transcriptional profiling in WT and the *ahk2,3* mutant. We identified differentially expressed genes (DEGs) between apical and basal internodes in both genotypes (Table S1). Considering the nature of the phenotype observed in *ahk2,3* (disturbed apical-basal developmental gradient), DEGs representing potential cytokinin targets and contributors to SCW formation were identified as (i) those genes showing a loss of WT apical-to-basal expression gradient in *ahk2,3* plants (3439 WT gradient-specific genes), or (ii) subset of intersecting genes whose apical-to-basal differential regulation differed by a factor ≥ 2 in amplitude between WT and *ahk2,3* (1499 out of 9050 intersecting genes revealing apical-to-basal gradient in both WT and *ahk2,3*; Fig. 2A, Table S2).

**Figure 2.**
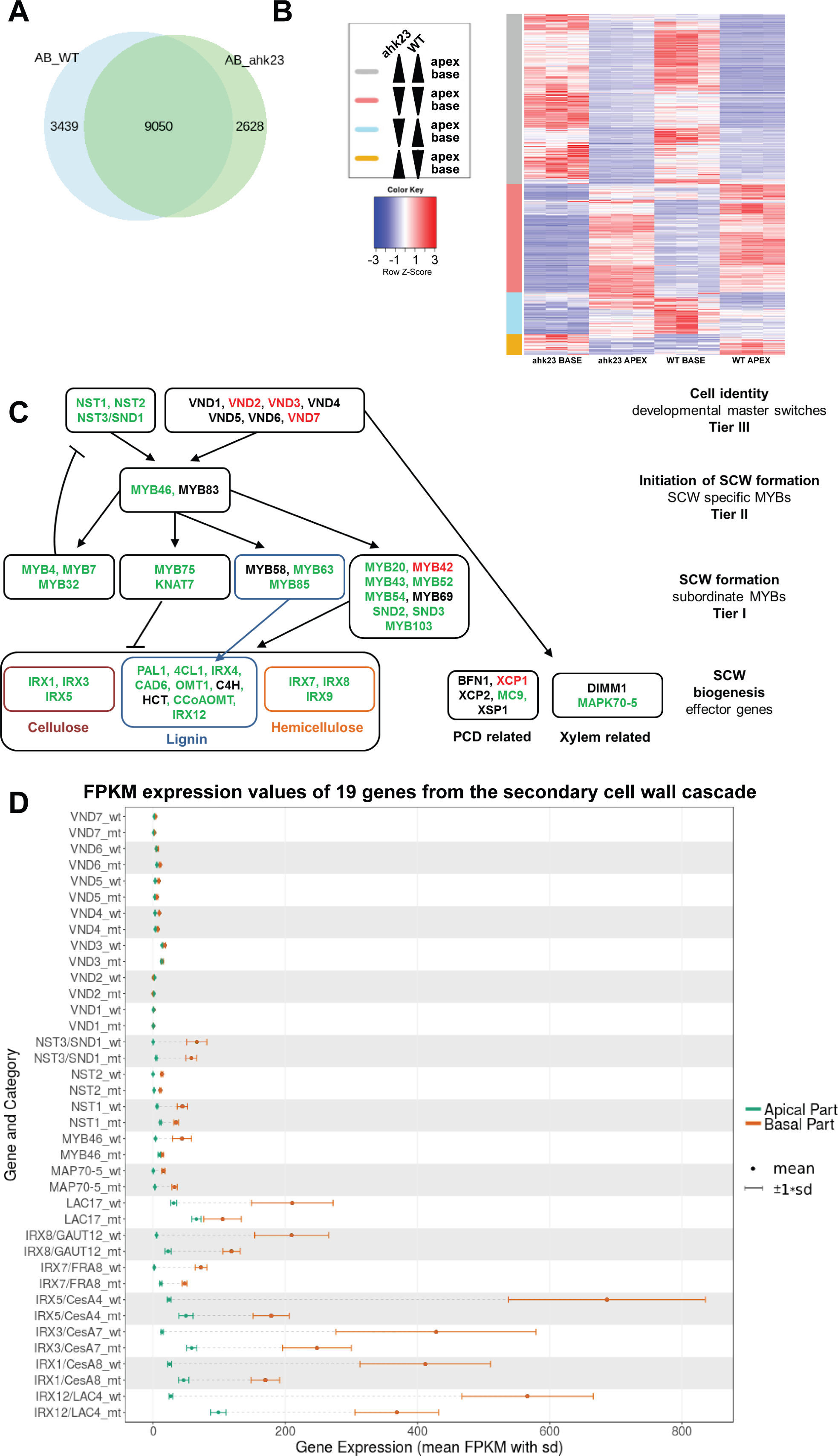
Cytokinins regulate the transcriptional cascade controlling the onset of SCW biosynthesis. **(A)** Venn diagram showing the overlap of differentially expressed genes between two experiments comparing apical versus basal internodes in WT Col-0 (AB_WT) and in *ahk2,3* mutant (AB_ahk23). The significance level was set at 0.05. **(B)** Heatmap of 1499 DEGs revealing the significant change in apical-to-basal expression in both WT and *ahk2,3*. At the same time, the apical-to-basal ratio of those genes differs by a factor ≥ 2 between WT and *ahk2,3*; for the list of the genes in individual subsets categorized according to the direction of the apical/basal difference in both genotypes (the color key on the left-hand side of the image) see Table S2. **(C)** Simplified scheme of the transcriptional cascade regulating SCW formation. Green and red indicate the upregulation or downregulation of the genes, respectively, in the apex of *ahk2,3* when compared to WT. **(D)** Comparison of the mean FPKM (Fragments Per Kilobase of exon per Million reads mapped) expression values of 19 genes from the SCW cascade in apical and basal internodes. Key: sd: standard deviation; mt: *ahk2,3* mutant; wt: Col-0 wild type.

In the case of the 3439 WT gradient-specific genes, the gene ontology (GO) analysis revealed ‘mRNA modification’ and ‘negative regulation of gene expression’ categories (Fig. S3A, Table S2) as significantly overrepresented categories. In comparison, the cell wall-related categories were frequently enriched among the 1499 intersecting genes, taken as a whole or separately for four possible gene subsets categorized according to the direction of the apical-basal difference in both genotypes (Fig. 2B, Fig. S3B-E, Table S2). Noteworthy, auxin signaling was strongly overrepresented in the subset showing higher basal than apical expression in both the WT and *ahk2,3* (Fig. S3C).

Based on published data, we constructed a multilevel regulatory cascade controlling SCW formation [Fig. 2C; (Didi et al., 2015; Hussey et al., 2013; Taylor-Teeples et al., 2015)]. At the top of the cascade, there are two groups of master regulators belonging to the group of NAC transcription factors: VASCULAR RELATED NAC-DOMAIN PROTEINs (VNDs) and NAC SECONDARY WALL THICKENING PROMOTING FACTORs (NSTs). While VNDs largely control SCW formation in TEs (Kubo et al., 2005; Zhou et al., 2014), NSTs are primarily responsible for SCW development in both xylary and interfascicular fibers (Mitsuda et al., 2007; Zhong et al., 2007). NSTs and VNDs regulate a battery of downstream transcription factors, which may act as either negative or positive regulators of SCW formation. These transcription factors orchestrate the expression of effector genes for cellulose, hemicellulose and lignin biosynthesis, as well as markers of xylem development, including genes for programed cell death and microtubule rearrangement during xylem formation (Fig. 2C).

The SCW-associated gene set (124 genes identified in the literature search, mostly members of the regulatory SCW transcriptional cascade, Table S3) was significantly enriched in the intersecting 9050 DFGs, showing apical-to-basal gradient in both WT and *ahk2,3* (Fig. 2A) when compared with all the protein coding genes detected in the whole dataset (Fisher’s exact test, *p-*value < 0.001). That is in the agreement with the apical-to-basal developmental gradient observed in the selected developmental stage of the *Arabidopsis* inflorescence stem (stage 1). In agreement with the observed phenotype, we detected an upregulation of most members of the SCW transcriptional cascade in apical internode of *ahk2,3* (highlighted in green in Fig. 2C). With few exceptions, the upregulation occurred mostly in the NST-regulated branch, including the upregulation of all *NSTs (NST1-NST3)*. Furthermore, the majority of the NST-regulated genes revealed similar changes in the apical-to-basal expression gradient (Fig. 2D, Table S3). In comparison to WT, in *ahk2,3* we observed a higher expression of *NSTs* and their downstream targets in apical (differentiating) portion of inflorescence stem, while lower expression of the NST-regulated genes was apparent in the basal part (fully differentiated cells). In contrast, most of the *VNDs* genes associated with TE formation showed similar apical-basal distribution in WT and *ahk2,3* plants, except some *VNDs* revealing slight changes in apical and basal parts of *ahk2,3* mutant (Fig. 2D). Thus, the cytokinin signaling deficiency in *ahk2,3* seems to preferentially activate the NST-regulated branch of the SCW regulatory cascade; however, the effect on VND-regulated SCW formation in vascular bundles cannot be excluded.

Taken together, these data demonstrate that the expression profile of the NST-controlled subset of SCW transcriptional cascade positively correlates with the developmental gradient observed along the apical-basal axis and suggest that cytokinin signaling controls the apical-basal gradient of SCW-related genes. Based on our data, cytokinins appear to act as negative regulators of *NST* master switches, thereby negatively regulating the entire downstream SCW transcriptional cascade in the apical portion of the inflorescence stem.

### Cytokinins downregulate the expression of *NSTs* to control secondary cell wall formation in apical internodes

To test if cytokinins act as negative regulator of *NSTs* expression, we assayed *NST3* expression in *proNST3:NST3-GUS* plants after the application of exogenous cytokinin (spraying the plants with 6-Benzylaminopurine (BAP) solution twice in 48 hours, Fig. 3A). In line with previous reports (Mitsuda et al., 2005), in the DMSO-treated controls we observed (weak) *NST3* expression predominantly in the interfascicular arcs; however, *NST3* activity was also detectable in the vascular bundles. In a good agreement with our transcriptional profiling data, there was an apparent increase in the *NST3* activity in the basal segments when compared to apical internodes, again both in the interfascicular regions and in the vascular bundles. In the plants treated with exogenous cytokinins (1 and 10 µM BAP), we observed a concentration-dependent inhibition of *NST3* expression (Fig. 3A).

**Figure 3.**
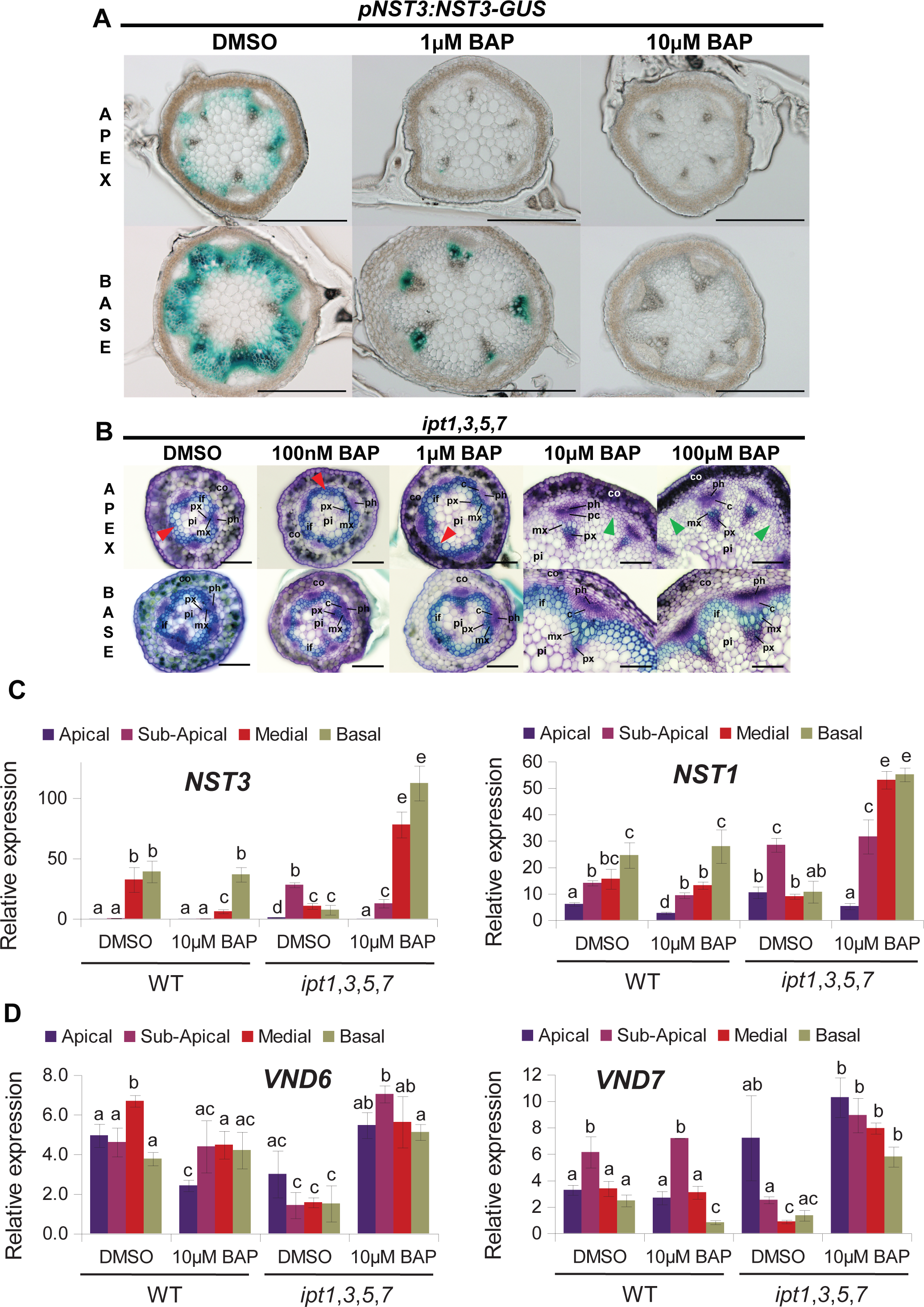
Cytokinins control the onset of SCW *via* the transcriptional regulation of *NSTs*. **(A)** Histochemical detection of *NST3* activity at the apex and base of the inflorescence stem in *pNST3:NST3-GUS* showing downregulation of *NST3* after 48 hours BAP treatment. Scale bars 200 µm. **(B)** Phenotype rescue of *ipt1,3,5,7* by exogenous BAP. Note the WT-like architecture of vascular bundles, upregulated procambial activity and absence of interfascicular arcs in apical internodia (green arrowheads) of plants treated with 10 µM and particularly 100 µM BAP. Red arrowheads point to the premature SCW formation. **(C-D)** RT-qPCR quantification of *NST3* and *NST1* **(C)**, and *VND6* and *VND7* (**D**) expression in apical, sub-apical, medial and basal internodes of inflorescence stems in control (DMSO)- and cytokinin (BAP)-treated WT Col-0 and *ipt1,3,5,7* mutant. Transcript levels were normalized to *UBQ10*. The relative *target gene*/*UBQ10* expression ratios are shown. Data are means ± SE from a representative experiment (n≥3). Different letters indicate significant differences at *P* < 0.05 based on a Tukey’s HSD test. The experiments were repeated 3 times with similar results.

Our genome-wide transcriptional profiling data suggest disturbed apical-basal gradient of *NSTs* in *ahk2, 3* background with higher levels in the apex, but lower at the base (Fig. 2D). This is implying cytokinins as negative regulators of *NSTs* specifically in the apex with possible consequences for *NST* activity in the older portions of the inflorescence stem. To corroborate the developmental importance of cytokinin-mediated *NST* regulations, we inspected the levels of *NST1* and *NST3* expression in the cytokinin biosynthesis-deficient *ipt1*,*3*,*5*,*7* quadruple mutant plants, both in the absence and presence of exogenous cytokinins (spraying the plants by BAP solution once a day for one week in the time interval from the onset of flowering until reaching stage 1 development). To get a better resolution, one cm portions of stems were collected at the apical, sub-apical, medial and basal internodes of the inflorescence (See Methods and Fig. S1A). In mock-treated *ipt1,3,5,7* plants, the precocious SCW formation (Fig. 3B) was associated with strongly disturbed *NSTs* expression gradients (assayed using RT-qPCR, Fig. 3C). When compared with the WT, *NST1* and *NST3* were strongly upregulated in the apical and sub-apical portions of *ipt1,3,5,7* inflorescence stems. On the other hand, we detected decreased levels of *NST3* in the medial internodia and both *NST1* and *NST3* were downregulated in the basal internodia of cytokinin-deficient line, confirming thus our genome-wide transcriptional profiling results in *ahk2,3* mutant (Fig. 2). Similar aberrant changes, i.e. upregulation in the apical internodia, but no change or even decrease in expression in the medial and basal internodia were observed for *IRX3* and *IRX8*, the downstream members of the NST-regulated SCW transcriptional cascade and markers of SCW-specific cellulose and hemicellulose biosynthesis (Fig. S4D). The expression of *VND6* and *VND7* involved in meta- and protoxylem development (Kubo et al., 2005) was downregulated in sub-apical, medial and basal portion of *ipt1,3,5,7* mutant (Fig. 3D).

While treatment with 0.1 and 1 µM BAP did not show any distinct phenotypic effects, application of 10 µM and 100 µM BAP inhibited precocious SCW formation in the apical part of the inflorescence stem of *ipt1,3,5,7* plants (Fig. 3B). Moreover, 10 µM BAP treatment partially rescued the growth defect (e.g. stem length) of *ipt1,3,5,7* mutant (Fig. S4A and S4B) and we observed a recovery of the stem diameter phenotype and number and architecture of vascular bundles (Fig. 3B), the traits previously demonstrated to be under the control of cytokinins (Hejatko et al., 2009; Matsumoto-Kitano et al., 2008; Nieminen et al., 2008). Importantly, the exogenous cytokinin treatment was able to rescue not only the SCW phenotype, but the *ipt1,3,5,7* mutant treated with 10 µM BAP plants also showed a WT-like apical-basal expression gradient of *NST1* and *NST3* revealing even higher levels in medial and basal internodia when compared to DMSO control (Fig. 3C). The expression of *IRX3* and *IRX8* in *ipt1,3,5,7* mutant was also rescued to WT-like pattern after treatment with 10 µM BAP; however, the rescue was not fully comparable to WT in case of *VND6* and *VND7* (Fig. S4A; Fig. 3D). Similarly to *NST1/NST3*, the expression of NST-regulated *IRX3/8* was exceeding the WT levels in BAP-treated *ipt1,3,5,7* (Fig. 3D and Fig. S4D).

To wrap it up, in line with RNAseq data discussed in the previous section, our results from cytokinin-deficient *ipt1,3,5,7* line support the idea that cytokinins act as regulators in the NAC TF-regulated SCW cascade. Proper levels of endogenous cytokinins seem to be necessary for preventing the precocious onset of SCW formation and apical/basal gradient of *NST1* and *NST3* expression. Interestingly, even the non-targeted application of exogenous cytokinins is able to rescue the developmental defects and recover the expression patterns of key TFs, particularly NST1/NST3, as well as their downstream targets.

### Cytokinin signaling controls expression of genes for NAC TFs along the apical/basal axis of inflorescence stem

To elucidate the role of AHK2/3-initiated cytokinin signaling in the regulation of SCW formation, we quantified the expression of selected NAC TFs and their downstream targets in mutants deficient in ARR1, ARR10 and ARR12, the type-B ARRs acting downstream of cytokinin-responsive AHKs and mediating dominantly cytokinin signaling (Argyros et al., 2008). We also used the *ahk2-2tk ahk3-3* double mutant, another allelic version of the *ahk2*,*3* (Higuchi et al., 2004) to confirm RNA-seq data performed on the *ahk2-1 ahk3-1* double mutant. Indeed, compared to Col-0 WT, we found *NST1*, *NST3* and *IRX3* strongly up-regulated in the apical, sub-apical and medial portions of the *ahk2,3* inflorescence stem (Fig. 4A, 4C and Fig. S5A). Similarly, *VND6* expression was higher in all stem portions of *ahk2,3* mutant compared to WT plants (Fig. 4E), while *VND7* was significantly up-regulated only in apical portion (Fig. 4G). Compared to that, the RT-qPCR analyses indicate that *VNDs* and *NSTs* expression profile in the *arr1*,*10*,*12* triple mutant is more similar to the *ipt1,3,5,7* than the *ahk2*,*3* background. In *arr1*,*10*,*12, NST3* was up-regulated in the apical and sub-apical portions but down-regulated in the basal part (Fig. 4B). Moreover, *NST1*, *VND7* and *IRX3* were slightly up-regulated in the apical or sub-apical portions but down-regulated in the medial and basal parts of *arr1*,*10*,*12* stem compared to Col-0 WT (Fig. 4D, 4H and Fig. S5B). *VND6* expression was repressed in both sup-apical and medial segments of *arr1*,*10*,*12* stem (Fig. 4F).

**Figure 4.**
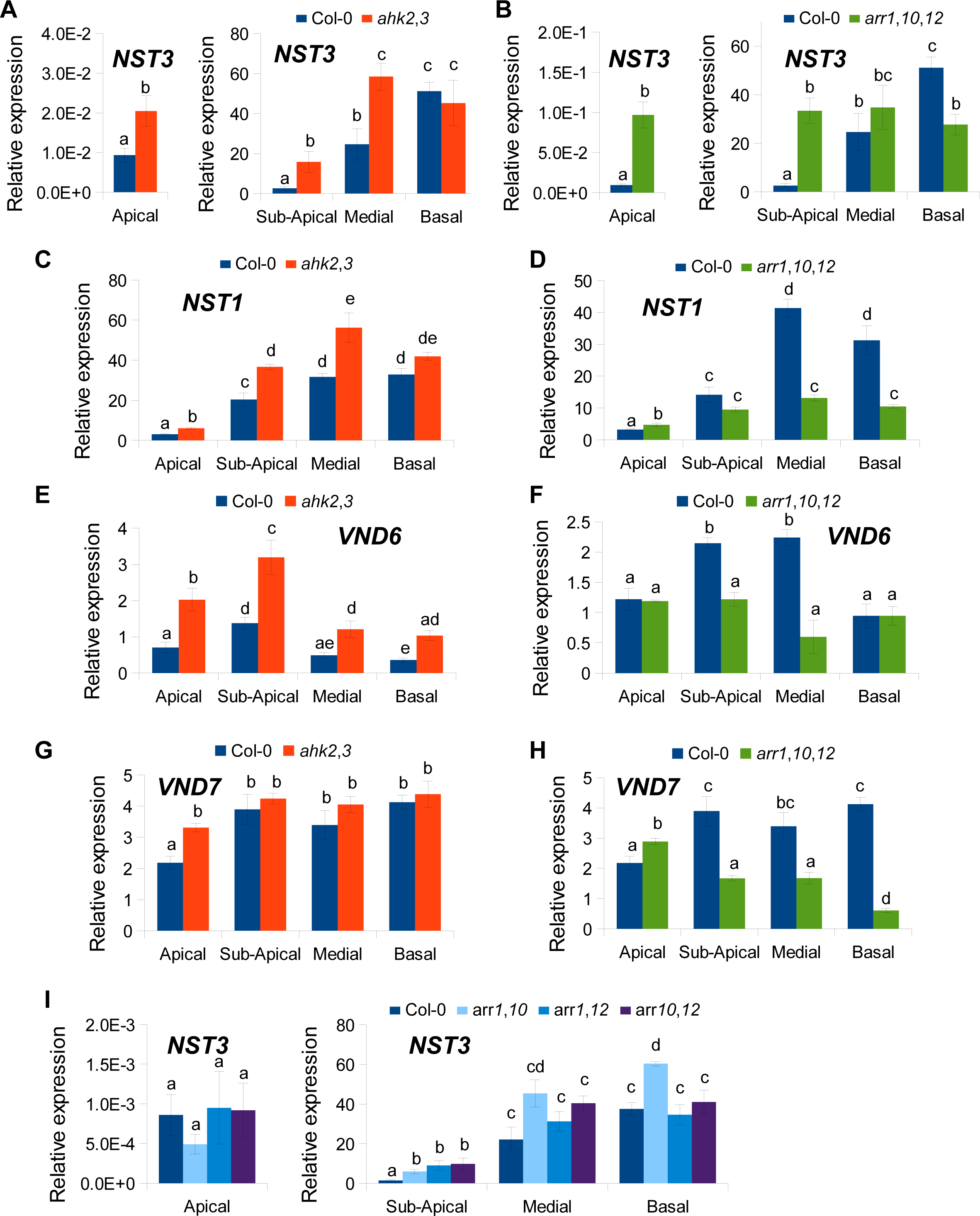
Defects in cytokinin signaling induce an up-regulation of *NST1*, *NST3* and *VND7* in the apical portion of stems. **(A**-**I)** RT-qPCR analysis of *NST3* **(A, B** and **I)**,*NST1* **(C** and **D)**, *VND6* **(E** and **F)** and *VND7* (**G** and **H**) expression in the apical, sub-apical, medial and basal portions of stems collected from WT (Col-0), *ahk2*,*3* double mutant **(A,C, E** and **G)**, *arr1*,*10*,*12* triple mutant **(B, D, F,** and **H)**, *arr1*,*10*, *arr1*,*12* and *arr10*,*12* double mutants **(I)**. Transcript levels were normalized to *UBQ10*. The relative *target gene*/*UBQ10* expression ratios are shown. Data are means ± SE from a representative experiment (n≥3). Different letters indicate significant differences at *P* < 0.05 based on a Tukey’s HSD test. The experiments were repeated 3 times with similar results.

As previously observed (Higuchi et al., 2004; Argyros et al., 2008; Hejatko, 2009), the *ahk2*,*3* and *arr1*,*10*,*12* mutants are affected in their growth and development (Fig. S6A). Compared to WT plants, *ahk2,3* and *arr1*,*10*,*12* showed a delay of about 5-7 days in the appearance of the first differentiated silique and a 30% to 40% reduction of the stem length respectively at the time of collection (Fig. S6B and C). Because a strong delay in flowering or a shorter stem length could influence SCW formation, we next analyzed the *arr1*,*10*, *arr1*,*12* and *arr10*,*12* double mutants which are less affected in their growth and development [(Argyros et al., 2008); Fig. S6]. Our phenotypic data indicated that in contrast to delay of flowering observed in triple *arr1,10,12* mutants, *arr1*,*10* and *arr1*,*12* double mutants exhibited an even slightly earlier appearance of the first differentiated silique (one to two days, respectively), when compared to WT (Fig. S6B). The *arr1*,*10* mutant also showed a slight but significant reduction in stem length of about 14% relative to WT (Fig. S6C). Importantly, the expression of *NST3* and *IRX3* was up-regulated in sub-apical portions of *arr1*,*10*, *arr1*,*12* and *arr10*,*12* double mutants compared to WT, while in the medial and basal portion *NST3* was significantly up-regulated only in the *arr1*,*10* mutant (Fig. 4I and Fig. S7B). This mutant also showed a slight but significant up-regulation of *NST1* expression in the sub-apical and apical portions (Fig. S7A).

To wrap it up, our results show a role of cytokinin-regulated MSP signaling in the regulation of expression pattern of NAC TFs along the inflorescence stem in Arabidopsis. We observed differences in the transcriptional regulation of the NAC transcription factors between the *ahk2*,*3* mutant (no or only small effect) and the *arr1*,*10*,*12* and *ipt1,3,5,7* mutants (repression) in the medial and basal parts of inflorescences, possibly suggesting more complex cytokinin-dependent regulation. Nevertheless, both the cytokinin receptor kinases AHK2 and AHK3 and the type-B ARRs ARR1, ARR10 and ARR12 likely repress SCW formation in the apical portion through the downregulation of *VND7*, *NST1*, *NST3* and *IRX3* expression. ARR1 (see also later in the text) and ARR10 may play a prominent role in repressing genes involved in SCW formation in the sub-apical and apical portions of the inflorescence stems.

### AHK2 and AHK3 act upstream of *NST1* and *NST3* in the control of SCW formation in interfascicular fibers but independently of *NST1/3* in the vascular bundles

To investigate the nature of genetic interaction between the AHK2- and AHK3-mediated cytokinin signaling and NAC TFs, the *ahk2-1* and *ahk3-1* mutations were introduced by crossing into the *nst1-1 nst3-1* (*nst1,3*) background, previously shown to be deficient in the SCW formation in the interfascicular arcs, but not in the TEs (Mitsuda et al., 2007). In quadruple *ahk2,3 nst1,3* mutants, we observed formation of smaller vascular bundles and reduced procambial activity, a phenotype typical for *ahk2,3*. Similarly to what we described for the *ahk2,3* in the text above (first section of Results, Fig. 1A), in both *ahk2,3* and *ahk2,3 nst1,3* we observed absence of differentiating TEs with primary CW and found metaxylem TEs with developed SCW to localize in a close proximity of procambium, suggesting accelerated xylem differentiation compared to WT. However, in contrast to the cytokinin (signaling) deficient lines, the presence of *nst1,3* resulted into a lack of SCW formation in the interfascicular fibers of *ahk2,3 nst1,3* inflorescence stem, both in the apical and basal internodia (Fig. 5). Interestingly, in the *ahk2,3 nst1* and *ahk2,3 nst3* triple mutants, a lack of SCW formation in the interfascicular fibers of inflorescence stem was also observed in the apical internode but not in the basal internode (Fig. S8). Nevertheless, in the basal portions of the *ahk2,3 nst1* triple mutant and to a lesser extend in the *ahk2,3 nst3* mutant, the interfascicular fibers showed a weaker toluidine blue staining compared to WT Col-0. This provides evidence that *nst1* and *nst3* are epistatic to *ahk2* and *ahk3* in the control of SCW initiation in the interfascicular arcs.

**Figure 5.**
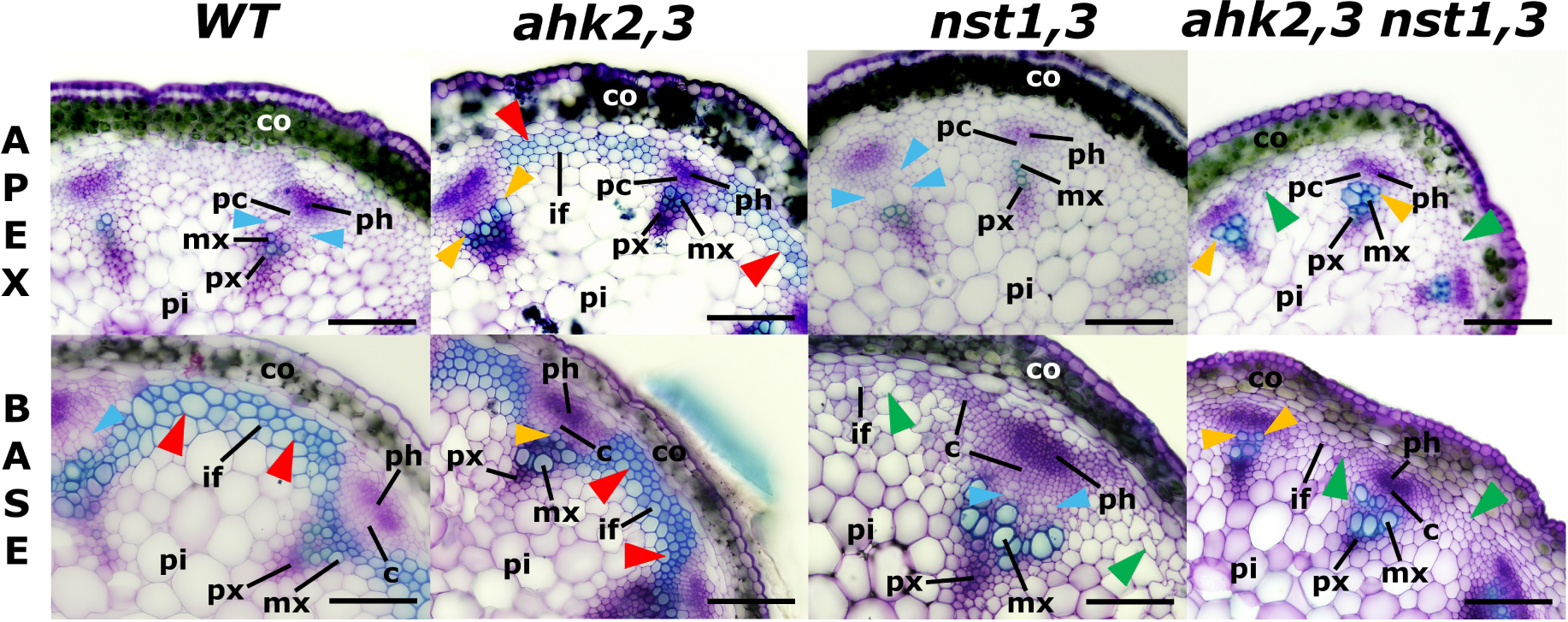
*AHK2* and *AHK3* act upstream of *NST1* and *NST3* in the control of SCW initiation. (**A**) Transverse sections through the inflorescence stem of WT *Col-0*, double mutants *ahk2-1 ahk3-1* (designated as *ahk2,3*) and *nst1-1 nst3-1 (nst1,3)* and quadruple mutant *ahk2-1 ahk3-1 nst1-1 nst3-1* (*ahk2,3 nst1,3*). Note the lack of the SCW formation in the interfascicular regions at the base of the inflorescence stem in *nst1,3* and both apex and base in *ahk2,3 nst1,3* (green arrowheads) when compared to WT and *ahk2,3* (red arrowheads). Also note the precocious SCW formation in differntiating TEs located very close to procambium in both *ahk2,3* and *ahk2,3 nst1,3* (orange arrowheads) compared to the absence of SCW in the differentiating TEs in WT and *nst1,3* (blue arrowheads). Smaller vascular bundles and reduced procambial activity is apparent in *ahk2,3* and *ahk2,3 nst1,3* as a result of attenuated cytokinin signaling. High resolution micrographs are provided allowing to zoom to even 800 % of the original image size to see the detailed VBs structure. Key: c – cambium; co – cortex; pi – pith; mx – metaxylem; px – protoxylem; pc – procambium; ph – phloem; if. interfascicular fibers. Scale bars: 50 μm

In summary, AHK2- and AHK3-regulated cytokinin signaling seems to control the onset of SCW formation *via NST1* and *NST3* in the interfascicular arcs. However, our data suggest existence of *NST1/3*-independent mechanism controlling SCW formation downstream of AHK2/3 in the xylem cells of vascular bundles.

### Cytokinin signaling is highly responsive in the apical and basal portions of the inflorescence stem

The aforementioned results suggest that cytokinins could be important in establishing the longitudinal expression pattern of NAC TFs. To analyze cytokinins distribution along the apical-basal axis, we measured the content of endogenous cytokinins along the inflorescence stem in WT and *ipt1,3,5,7* plants (Fig. 6). In WT apical internodes, we observed slightly higher levels of both *trans*-zeatin (*t*Z) and *N*^6^-(Δ^2^-isopentenyl)adenine (iP), considered the dominant active cytokinins in plants, when compared to medial internodes. The effect was even more pronounced if the *t*Z and iP ribosides (*t*ZR and iPR, respectively), proposed to act dominantly as activatable transport cytokinin forms, were included. On the other hand, we also observed a slight increase in both *t*Z and iP cytokinins in the basal segments relative to medial internodes, resulting into a “V-shaped” distribution of endogenous cytokinins along the apical-basal axis in the *Arabidopsis* inflorescence stem.

**Figure 6.**
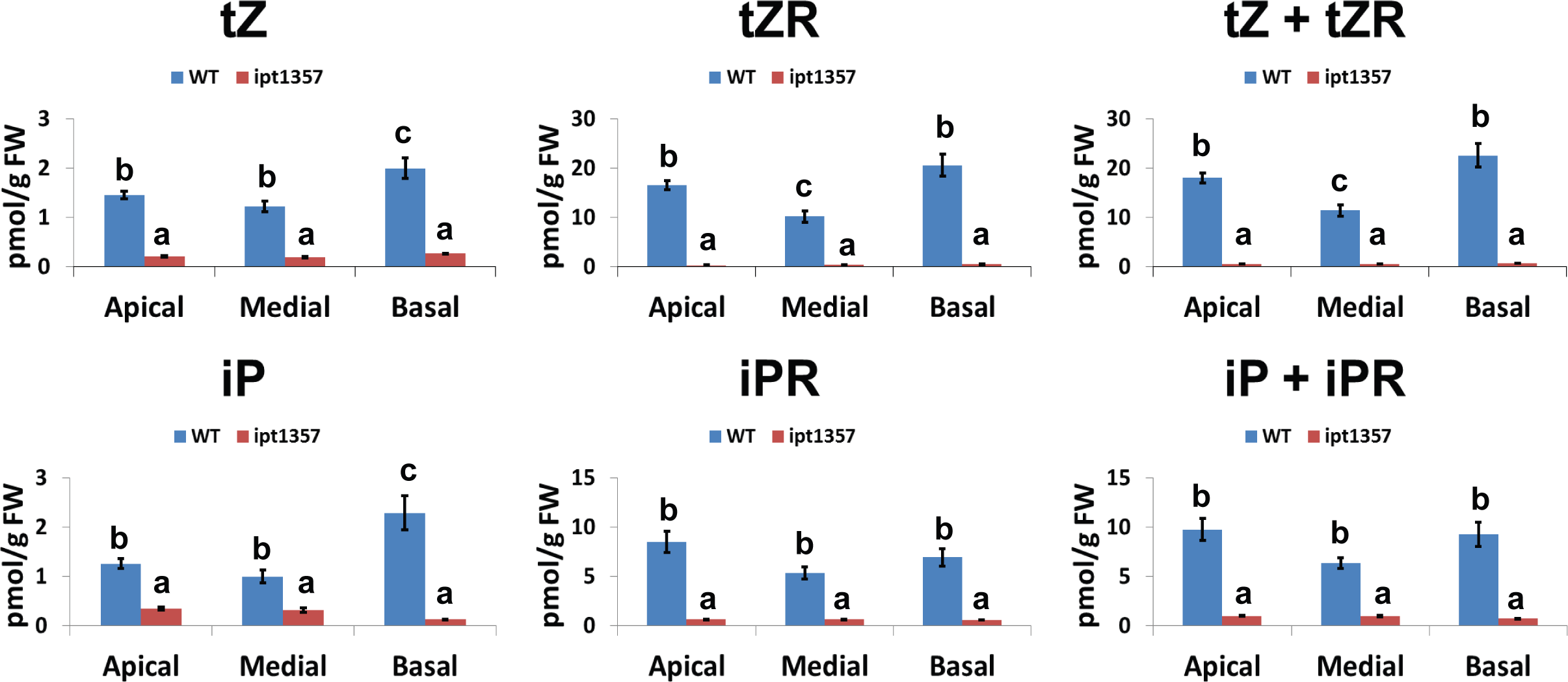
Endogenous cytokinin distribution reveals only small variability along the apical-basal axis of the inflorescence stem. Measurement of endogenous cytokinin species tZ, trans-zeatin; tZR, tZ riboside; iP, N^6^-(Δ^2^-isopentenyl)adenine; iPR, iP riboside in the apical, medial and basal internodes of Col-0 WT and *ipt1*,*3*,*5*,*7* mutant. Data are means ± SE from a representative experiment (n≥4). Different letters indicate significant differences at *P* < 0.05 based on a Tukey’s HSD test.

When compared to WT, the levels of *t*Z and iP in *ipt1,3,5,7* were strongly downregulated in all (apical, medial and basal) internodes (Fig. 6). In good agreement with previously reported data (Miyawaki et al., 2006) the opposite effect, *i.e.* the upregulation of *cis-*zeatin (*c*Z), was observed in *ipt1,3,5,7* when compared to WT (Fig. S9). This implies the activation of a compensatory mechanism, possibly mediated by the *AtIPT2* and/or *AtIPT9*, which are proposed to be responsible for *c*Z biosynthesis in the *AtIPT*-deficient line (Miyawaki et al., 2006). However, *c*Z was shown to be much less active in most cytokinins bioassays, and its role in the cytokinin-mediated regulation of plant development remains uncertain (Gajdosova et al., 2011; Hosek et al., 2019; Schafer et al., 2015).

The V-shaped endogenous cytokinin distribution was also partially reflected in the activity of cytokinin signaling, as could be seen from the expression of cytokinin–responsive type-A ARRs (*ARR5, ARR7* and *ARR15*). Although the differences observed in the WT were rather small and significantly higher expression in apical and basal internodia compared to medial ones was detectable only for *ARR15* (and partially also in case of *ARR7)*, the higher expression in apical and basal internodia compared to subapical and/or medial internodia of all assayed type-A ARRs (*ARR5, ARR7* and *ARR15*) was seen in BAP-treated *ipt1,3,5,7* (Fig. S4E-F), suggesting higher cytokinin responsiveness particularly in apical and basal internodia. *ARR5, ARR7* and *ARR15* expression was strongly decreased in all internodia of DMSO-treated *ipt1,3,5,7*, pointing to the impaired cytokinin signaling in cytokinin-deficient lines (Fig. S4E-F).

The expression of *ARR5* and *ARR7* was also analyzed in the stems of *ahk2,3* double mutant and type-B ARR triple and double mutants to verify that cytokinin signaling is effectively repressed in these mutants (Fig. S5C-F and S7C, D). With the exception of *ARR5* in the *ahk2,3* mutant, our RT-qPCR results indicate that cytokinin signaling is generally down-regulated in all stem portions (apical, sub-apical, medial and basal) of the of *ahk2,3*, *arr1*,*10*,*12*, *arr1*,*10* and *arr1*,*12* mutants, but not in *arr10,12*. Therefore, ARR1 is probably the major type-B ARR transcription factor regulating cytokinin response in the inflorescence stem.

To understand the possible mechanism of differential cytokinin signaling responsiveness along the longitudinal axis of the inflorescence stem, we analyzed the expression of type-B ARRs *ARR1, ARR2, ARR10, ARR11* and *ARR12* in the individual sections of inflorescence stem in Col-0 WT plants. With the exception of ARR10 and ARR11, revealing higher expression in basal/medial internodia compared to apical/subapical ones, we did not observe strong differences in the expression of type-B ARRs along the apical/basal axis of the inflorescence stem (Fig. S10).

In conclusion, the results of our measurements indicate that higher levels of endogenous cytokinins are correlated with elevated cytokinin signaling activity in the apical and basal portions of Arabidopsis inflorescence stem. Together with recovery of the V-shape pattern of cytokinin signaling even after equal application of exogenous cytokinin to the entire flowering *ipt1,3,5,7* plants, it is implying spatially-specific responsiveness of cytokinin signaling pathway. However, rather uniform activity of genes for type-B ARRs cannot explain this type of spatial-specific cytokinin response.

### Cytokinins control functional properties of water conducting elements

To assess the functional properties of TEs along the apical-basal axis in the WT inflorescence stem, we measured hydraulic conductivity in the individual inflorescence internodes. The hydraulic conductivity is a function of total TE area that depends on both the number and diameter of functional TEs. And, it has been shown that water conductivity increases exponentially (to the fourth power) with increasing vessel diameter (Tyree et al., 1994). Our analysis showed a remarkable increase in the hydraulic conductivity of the basal segments compared to apical internodes (Fig. 7A). We visualized functional TEs using the recently developed protocol of Jupa et al. (2015). Apparently, increases in both TE number (particularly of metaxylem type) and TE diameter contribute to the substantive increase in hydraulic conductivity along the apical-basal axis of WT inflorescence stems (Fig. 7B, Fig. S11).

**Figure 7.**
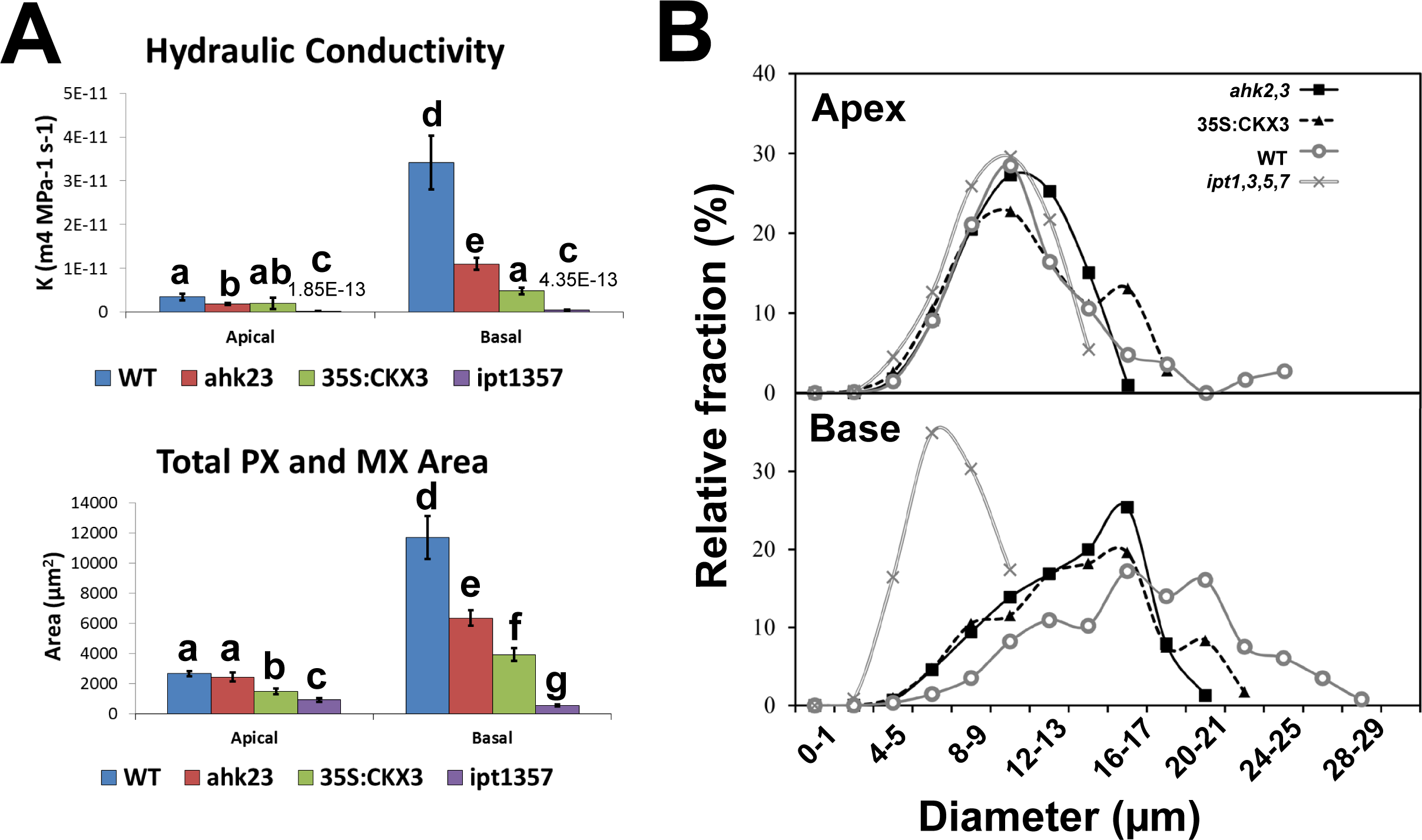
Cytokinin (signaling) deficiency impairs hydraulic conductivity of functional tracheary elements. **(A)** Hydraulic conductivity and total xylem lumen area in the inflorescence stems of WT, and cytokinin (signaling) deficient lines (±SE); values are shown in the chart for the (very low) hydraulic conductivity of *ipt1,3,5,7* line (±SE). Data are means ± SE from a representative experiment (n≥6). Different letters indicate significant differences at *P* < 0.05 based on a Tukey’s HSD test. **(B)** The distribution of the individual classes of proto- and metaxylem TEs clustered according to their diameter in the apical and basal internodes of WT and cytokinin (signaling) deficient lines.

To assess the possible influence of premature SCW formation on TE properties in cytokinin (signaling) deficient lines, we assayed hydraulic conductivity of functional TEs in the inflorescence stem of *ahk2,3*, *p35S:CKX3* and *ipt1,3,5,7* lines. Relative to WT, we found that all mutant and transgenic lines tested demonstrated decreases in the total proto- and metaxylem area with a strongly impaired hydraulic conductivity in the basal internode, and in most of these lines (2 out of 3), this was also observed in the apical internode (Fig. 7A, Fig. S11A). The strongest effect on the water conductivity gradient (loss of the statistically significant difference between the apical and basal internode) was apparent in the quadruple mutant *ipt1,3,5,7*. This is in line with a strong upregulation of SCW formation in the apical internode of the quadruple *atipt* mutant, leading to the almost complete absence of developmental gradient in SCW formation (Fig. 1). Interestingly, in the *ahk2,3* mutant, the total TE number (both proto- and metaxylem type) was comparable to that observed in WT (with metaxylem TEs being even upregulated compared to WT; Fig. S11A). However, the average diameter of functional TEs was decreased, leading to a decline in the total TE area and reduction in the hydraulic conductivity (Fig. 7A). The same trend, *i.e.* decrease in the diameter of functional TEs, was observed in *35S:CKX3* and particularly in the *ipt1,3,5,7.* In both of these lines, the drop in the cell number also accounted for the decline in the total TE area (Fig. 7A, Fig. S11A). Even more importantly, the absence of fraction of largest TEs, being responsible for the majority of hydraulic conductance in the WT (Fig. S11B), seems to be the main factor leading to the decreased hydraulic conductivity in all cytokinin (signaling) deficient lines (Fig. 7B).

Together, the data indicates that premature SCW formation or alteration in *VNDs* expression, induced by deficiency in particularly AHK2- and AHK3-mediated cytokinin signaling or endogenous cytokinin levels, can be associated with the decrease in a diameter of functional TEs, resulting in a strongly impaired water conductance of shoot vasculature.

### Ectopic overexpression of *NST1* mimics the effects of cytokinin deficiency on secondary cell wall formation and hydraulic conductivity

To inspect the possible effect of (cytokinin-mediated) misregulation of NAC TFs on SCW formation and the functional properties of TEs, we inspected SCW formation and water conductivity of *nst1,3* and *p35S:NST1* plants. In agreement with previous reports, we observed that *nst1,3* plants display absence of interfascicular arcs, but are unaffected in the onset of SCW in TEs (Figs. 5, 8A). Accordingly, we observed a WT-like ability to conduct water in both apical and basal internodes of *nst1,3* plants (Fig. 8B). In contrast, *p35S:NST1* mimics the enhanced SCW formation both in the vascular bundles and interfascicular arcs. In the VBs, the overexpression of *NST1* mimics the phenotype observed in the cytokinin (signaling) deficient lines, i.e. absence of differentiating TEs surrounded with primary CW and formation of cells with SCW in the close vicinity of (pro)cambium or cambial cells, even facing the phloem (Fig. 8A). That also associated with a decreased hydraulic conductivity in basal internodes relative to WT (Fig. 8B). Thus, similarly to the strong phenotype of *ipt1,3,5,7*, *p35S:NST1* also lacks an apical-basal gradient in hydraulic conductivity.

**Figure 8.**
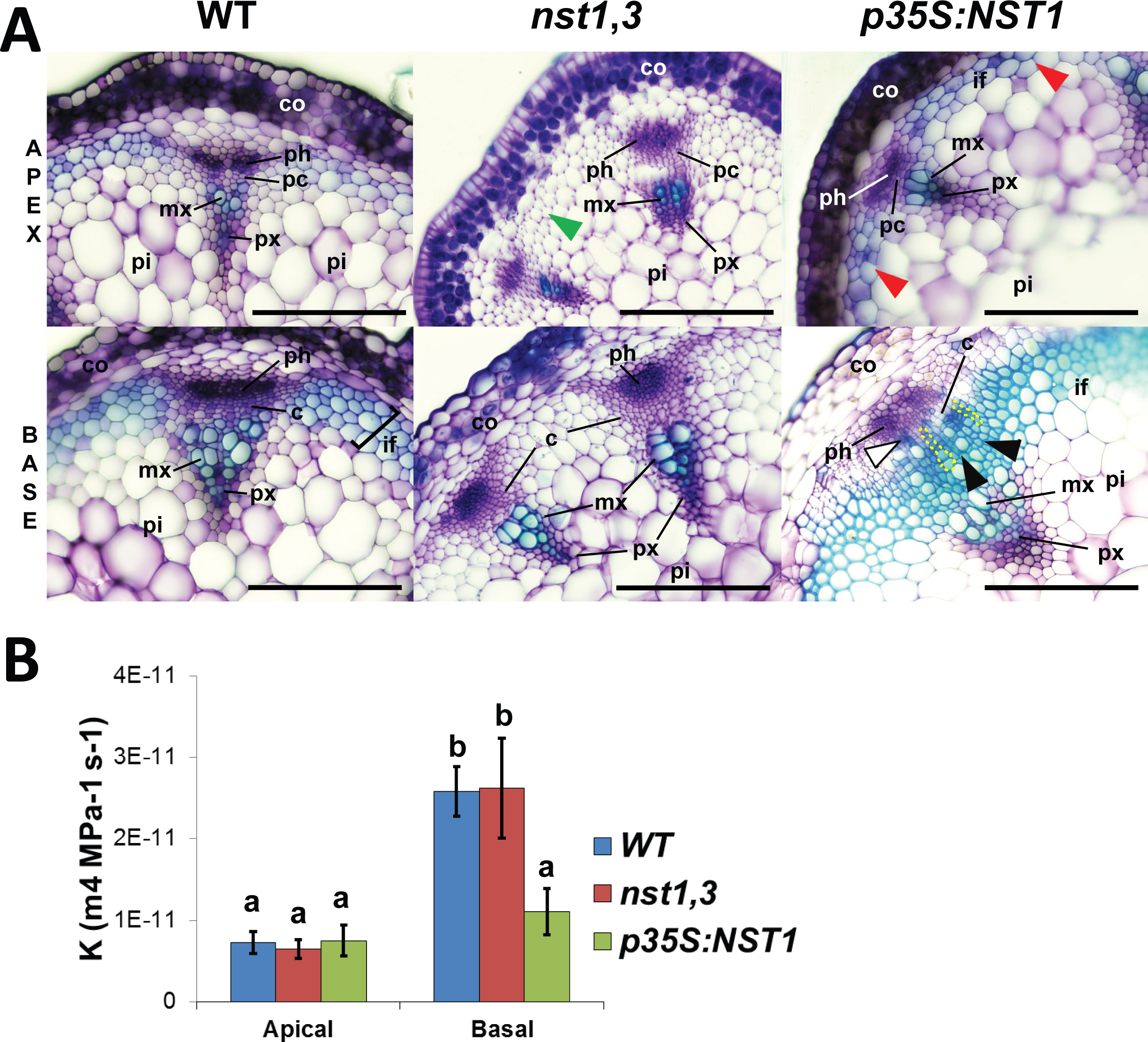
The ectopic overexpression of *NST1* mimics cytokinin (signaling) deficiency effects on SCW formation and hydraulic conductivity. **(A)** Toluidine blue-stained transverse sections of living WT Col-0*, nst1-1 nst3-1* and *p35S:NST1* inflorescence stems. Note the premature formation of SCW in the interfascicular xylem fibers/parenchyma in the apical internodes of *p35S:NST1* plants, while its absence in *nst1,3* (red and green arrowheads, respectively), in the latter case even at the base of the stem. Note also stronger toluidine blue staining at the base of the *p35S:NST1* stems and formation of radial arrays of xylary procambial-like cells (black arrowheads or yellow dashed line) revealing precocious SCW formation and formation of SCW in cambial cells even facing the phloem (white arrowheads). High resolution micrographs are provided allowing to zoom to even 800 % of the original image size to see the detailed VBs structure. Abbreviations as in 1A; scale bars: 50 µm. **(B)** Hydraulic conductivity in apical and basal internodes of WT Col-0*, nst1,3* and *p35S:NST1* inflorescence stems (±SE). Data are means ± SE from a representative experiment (n≥12). Different letters indicate significant differences at *P* < 0.05 based on a Tukey’s HSD test.

In summary, these data show that elevated expression of *NST1* partially phenocopies the developmental aberrations observed in both the vascular bundles and interfascicular arcs in cytokinin (signaling) deficient lines.

## Discussion

### Cytokinins control NAC TF-regulated SCW formation

We observed clear developmental gradient along the apical/basal axis of the inflorescence stem in *Arabidopsis*, associated with gradient in the expression (low in apex, high at the base) of genes for NST1/3 master regulators controlling SCW transcriptional cascade. Impaired cytokinin signaling or deficiencies in endogenous cytokinins results in perturbed xylem and interfascicular arcs development characterized by premature SCW formation and alterations in the apical/basal expression pattern of genes encoding NAC TFs as well as their downstream targets acting in the SCW-inducing transcriptional cascade. The observed differences in the expression of *NST1/3* and *VND6/7* in the apical and/or subapical portion of the inflorescence stem (low in WT but high in cytokinin signaling deficient *ahk2,3* line) together with high activity of cytokinin signaling in apex suggest that cytokinin-mediated downregulation of genes for NAC TF in the apical/subapical internodia is an important part of a mechanism allowing formation of proper developmental gradient along the apical-basal axis of the inflorescence stem.

Although the detailed mechanism remains to be clarified, our data suggest that AHK2 and AHK3 downregulate the expression of the genes for NAC TF *NST1/3* and *VND6/7* in the apical and *NST1/3* and *VND6* in the subapical portions of the inflorescence stem. Further, we demonstrate that type-B ARRs ARR1, ARR10 and ARR12 act redundantly in repressing the expression of *NST1* and *NST3* in apical portions and that ARR1 and ARR10 may play a prominent role in this process. The analysis of *ARR5* and *ARR7* expression in *arr1*,*10*, *arr1*,*12* and *arr10*,*12* double mutants also indicated that ARR1 is the main type-B ARR activating cytokinin signaling along the inflorescence stems.

The developmental gradient can be seen not only along the apical-basal axis, but also at the level of individual vascular bundles. In the WT, the differentiating (expanding) metaxylem TEs located proximally to procambial cells are still surrounded by primary cell wall and the SCW is formed in the developmentally older cells located more centrally to the inflorescence stem. Compared to that, in the cytokinin (signaling) deficient lines, this developmental gradient is much steeper and vast majority of cells morphologically distinguishable as future metaxylem TEs develops SCW although still located in the close vicinity of procambium. Thus, it seems that AHK2- and AHK3-mediated cytokinin signaling delays precocious SCW formation not only in the interfascicular arcs, but also in the differentiating TEs within vascular bundles.

The differences in the expression of the genes for NAC TFs we have observed between *ahk2,3* on one side and *arr1*,*10*,*12* and *ipt1,3,5,7* on the other one seem to be the result of strong developmental aberrations we observed in the inflorescence stem of *arr1*,*10*,*12* and *ipt1,3,5,7*, confirming the previously described role of cytokinin signaling in both primary and secondary growth (Hejatko et al., 2009; Matsumoto-Kitano et al., 2008; Nieminen et al., 2008). The weak upregulation of *VND7* and no upregulation of *VND6* in the apical internodia in case of *arr1,10,12* and *ipt1,3,5,7* can be a result of strongly impaired primary growth (highly reduced procambial activity) resulting in small vascular bundles and in case of *ipt1,3,5,7* also decreased number of vascular bundles, thus leading to reduction in a number of SCW forming cells. Besides the differences in the expression of *VND6/7*, there is also absence of *NST1* and *NST 3* downregulation in medial and basal internodia in *ahk2,3* line compared to *ipt1,3,5,7* and *arr1,10,12*. This seems to be the result of differential effects of cytokinin signaling, being much more disturbed in *ipt1,3,5,7* and *arr1,10,12* compared to *ahk2,3*. This can be well demonstrated on the expression pattern of type-A ARRs, particularly *ARR5*, being strongly downregulated in all internodia of both *arr1,10,12* and *ipt1,3,5,7*, but nearly not affected in *ahk2,3*. On the other hand, *ARR7* is downregulated to a comparable extent in apical and subapical internodia in both *ahk2,3* and *arr1,10,12*, but it is much more downregulated in medial and basal internodia of *arr1,10,12* than in the corresponding stem fragments of *ahk2,3*. Thus, the only remaining cytokinin sensor AHK4 is apparently able to mediate partial cytokinin signaling in *ahk2,3*, however, with some spatial specificity (medial and basal internodia) and revealing preference for the individual proteins within the MSP signaling (both type-B and type-A ARRs, as can be judged on the dominant role of ARR1 in mediating the AHK2- and AHK3-dependent cytokinin signaling in the inflorescence stem). This is in line with previous findings, suggesting certain level of specificity of individual cytokinin sensors (Higuchi et al., 2004; Nishimura et al., 2004; Riefler et al., 2006). The AHK4-mediated cytokinin signaling in medial/basal internodia may be possibly playing a positive role in the secondary growth (developmental switch of procambium to cambium), as recently demonstrated in the root (Ye et al., 2021), being responsible for the formation of SCW-containing (and thus NAC TFs expressing) cells at the base of the Arabidopsis inflorescence stem. Furthermore, because of precocious and excessive SCW production in strong cytokinin mutants, unnecessary SCW synthesis can be repressed or at least not activated as an adaptive mechanism in later stages of inflorescence development in cytokinin-deficient lines. The apparent increase in SCW deposition at the base of inflorescence of strong cytokinin mutants supports this hypothesis. In this scenario, negative feedback regulation of *NSTs* master switch by downstream MYB transcription factors is possible (Wang et al., 2011), thus explaining the downregulation of *NACs* in the medial and basal internodia of *arr1,10,12* and *ipt1,3,5,7*.

Cytokinins and AHK2- and AHK3-mediated cytokinin signaling is required for the (pro)cambium activity and radial growth (Hejatko et al., 2009; Nieminen et al., 2008). Consequently, the defects in primary growth observed in *ahk2*,*3* and particularly *ipt1,3,5,7*, and *arr1*,*10*,*12* mutants may induce compensatory mechanisms leading to early SCW formation. However, the number of protoxylem and metaxylem TEs is comparable in the WT and *ahk2,3*, suggesting the defect in the primary growth (proliferative procambium activity) is not that strong and can be partially compensated by the upregulation of metaxylem TEs differentiation. Furthermore, even though the *arr1*,*10*, *arr1*,*12* and *arr10*,*12* double mutants are less affected in their development (e.g. these lines do show stem length and flowering time comparable with WT Col-0), there is still significant upregulation of *NST3* as well as its downstream target *IRX3* detectable in the sub-apical stem fragment in the double *arr* mutants, suggesting that *NST*-regulated SCW is under negative control of cytokinin signaling in the early stages of primary growth of the inflorescence stem.

In spite of key importance of cytokinin signaling and endogenous cytokinins demonstrated in our study, cytokinins may cooperate with other factor(s), both positive and negative regulators of SCW formation, in achieving proper timing of SCW thickening. The results of our GO analysis together with the extensively described role of auxin in the control of xylem differentiation (Ruzicka et al., 2015) and xylem fiber and TEs expansion (Nilsson et al., 2008) makes auxin a likely candidate. Noteworthy, the overexpression of auxin-inducible *AtHB8* results in a phenotype resembling that of a cytokinin deficiency [precocious differentiation/SCW formation of interfascicular arcs; (Baima et al., 2001)], implying auxin as a potential positive regulator of SCW formation. A strong mutually negative interaction between cytokinins and abscisic acid (ABA) signaling was described (Skalak et al., 2021; Zubo and Schaller, 2020). Accordingly, ABA was shown to act as positive regulator of SCW formation in Arabidopsis and *VND7* and *NST1/3* were up-regulated by elements of ABA signaling pathway (Campbell et al., 2018; Ramachandran et al., 2021). Furthermore, SnRK2 kinases acting in ABA signaling were shown to phosphorylate NST1, allowing to control the NST1-regulated genes acting in the SCW formation (Liu et al., 2021). Thus, ABA could be another partner of cytokinins in the control of SCW formation.

### Cytokinin-regulated onset of SCW formation impacts on TE hydraulic conductivity

Our results indicate that precocious SCW in cytokinin insufficient lines leads to drop in hydraulic conductivity due to formation of smaller TEs. Importantly, thanks to our previously introduced approach (Jupa et al., 2015), we were able to correlate the total inflorescence stem water conductivity with the size of only functional TEs, thus avoiding possible bias due to including the size of non-functional TE/TE-like cells. Although NSTs were shown to induce SCW formation largely in the interfascicular fibers (Mitsuda et al., 2007; Zhong et al., 2007), both *NST1* and *NST3* were found to be expressed not only in the interfascicular regions, but also in the xylem cells differentiating into vascular vessels (Mitsuda et al., 2007). Furthermore, it was observed that *NST3* overexpression induces a slight increase in thickness of SCW in TEs but a decreased SCW in xylary and interfascicular fibers (Zhong et al., 2006, Ko et al., 2007) suggesting that regulation of SCW formation in both interfascicular and xylary fibers need a proper NST3 dosage. This data is in a good accordance with our findings that the premature formation of SCWs in the differentiating TEs observed in *ahk2,3* but also in other cytokinin (signaling) deficient lines can be partially phenocopied by *NST1* overexpression, suggesting that misregulation of *NST1/3* is able to affect not only interfascicular arcs, but also TEs differentiation. However, in terms of the TE size, there will be probably relevant also upregulation of *VND6* and *VND7,* previously shown to be responsible for the SCW formation in metaxylem TEs and protoxylem, respectively (Kubo et al., 2005). This seems to be the case particularly in the absence of functional *NST1/3* in *ahk2,3 nst1,3* quadruple mutant line, where the upregulation of *VND6* and/or *VND7* is probably responsible for the early SCW formation in the differentiating metaxylem TEs.

### Conclusions and future outlines

Based on our findings, we propose that high activity of AHK2- and AHK3-mediated cytokinin signaling in the apical portion of the inflorescence stem prevents premature SCW formation in vascular bundles as well as interfascicular regions, thus ensuring the proper timing of both xylem and interfascicular fiber differentiation. In the absence of endogenous cytokinins and/or attenuated cytokinin signaling, the ectopic upregulation of genes for NAC TFs initiates precocious formation of SCWs, before the cell expansion of differentiating TEs is complete. The premature formation of a rigid SCW prevents TEs from reaching normal diameters, thus impairing their hydraulic conductivity (Fig. S12).

Importance of cell wall properties in the developmental regulations in plants is emerging (Braybrook and Jonsson, 2016; Didi et al., 2015; Chebli and Geitmann, 2017; Sassi and Traas, 2015; Trinh et al., 2021). Our results imply that the molecular machinery controlling the onset of SCW formation is an important target of hormonal regulations, strongly affecting progression of differentiation and consequently functional properties of xylem cells. Our study clearly indicates that cytokinin signaling pathway plays a positive role in controlling long-range water transport in plants. This further supports the importance of MSP signaling in mediating the plant growth and highlights the individual members of the pathway as valuable targets of molecular-assisted breeding and/or synthetic biology approaches in the efforts to improve plant biomass formation.

## Materials and methods

### Plant materials and growth conditions

The double mutants *ahk2-2tk ahk3-3*, *ahk2-1 ahk3-1* (*ahk2*,*3*), *arr1-3 arr10-5* (*arr1*,*10*), *arr1-3 arr12-1* (*arr1*,*12*), *arr10-5 arr12-1* (*arr10*,*12*), the triple mutant *arr1-3 arr10-5 arr12-1* (*arr1*,*10*,*12*), the triple *atipt3-2 atipt5-1 atipt7-1* (*ipt3*,*5*,*7*), the quadruple *atipt1-1 atipt3-2 atipt5-1 atipt7-1* (*ipt1*,*3*,*5*,*7*) mutant, the quintuple mutant *ahp1-1 ahp2-1 ahp3 ahp4 ahp5-1* (*ahp1*,*2*,*3*,*4*,*5*), the *p35S:CKX2* and *p35S:CKX3* overexpressing lines and the *proNST3:NST3-GUS* line are in *Arabidopsis* Col-0 background and were previously described (Werner et al., 2001; Nishimura et al., 2004; Higuchi et al., 2004; Argyros et al., 2008; Hutchison et al., 2006; Miyawaki et al., 2006; Zhong et al., 2006). Plants were grown on soil (compost [TS-3; Klasmann-Deilmann], perlite and sand in the ratio of 12:3:4 respectively) in a growth chamber under long-day conditions (16 h light at 21°C/8 h dark at 19°C), at 60% humidity and illuminated with fluorescent tubes at 100 µmol m^−2^ s^−1^ light intensity. Stem samples were collected when the plants reach the developmental stage 1 of the first elongated silique (Altamura et al., 2001).

### Plant transversal sectioning

50 µm-thick cross sections were prepared from apical and basal internodes of *Arabidopsis* inflorescence stems at stage 1 (Altamura et al., 2001) by a vibrating blade microtome (Leica VT1200 S), stained with a 0.05% (w/v) solution of toluidine blue in water for 1 min and rinsed in distilled water three-times for 30 s. The water-mounted native cross sections were observed using microscope [Olympus BX61; (digital camera – Olympus DP70)]. Pictures of each individual vascular bundle were photographed at 4x, 10x, 20x or 40× magnification.

### BAP treatment and GUS staining

*proNST3:NST3-GUS* in Col-0 background were grown under the conditions as described above. Plants were sprayed once per day for 48 hours with 1 µM, 10 µM, 100 µM BAP or mock (0.1 % DMSO, Sigma). Small pieces (approximately 0.5 cm) of apical and basal portion of the inflorescence stem were cut and incubated in GUS staining buffer (0.1% Triton X-100, 1 mM X-GlcA sodium trihydrate, 20% methanol and 0.5 mM potassium ferricyanide and 0.5 mM potassium ferocyanide) for 24 hours under vacuum in darkness and at room temperature. Stained parts were analyzed as is described in plant transversal sectioning section.

### Illumina library construction and sequencing

Total RNA was extracted and used for library construction and sequenced using Illumina TrueSeq protocols by <u>GATC Biotech AG</u>, Constance, Germany. After removal of low-quality reads, >30 million mapped reads were retained for further analysis from each sample. In total, the expression of ∼26,900 distinct protein coding genes was detected.

### Sequence alignment

The adapters were removed using Cutadapt (Martin, 2011). Reads were mapped to the reference *A. thaliana* genome (TAIR10) using TopHat 2.0.8 (Kim et al., 2013) with the following parameters: minimum intron length, 20 bp; maximum intron length, 4 000 bp; only reads across junctions indicated in the supplied GFF (TAIR10) were utilized; all other parameters were set to default settings).

### Differential gene expression analysis

Cuffdiff 2.2.1 software (Trapnell et al., 2013) was used with default settings with a false discovery rate of 0.05. The most biologically relevant genes for SCW development were identified as differentially expressed genes showing the same or opposite direction of change between apical and basal internodes and with a significantly different amplitude of change (log2 fold changes ≥ 1.1 (log2 fold change_ahk2 ahk3_ – log2 fold change_WT_| >= 1.1)).

### Gene set enrichment analysis

We performed a gene enrichment analysis using GOrilla software (Eden et al., 2009) with a FDR value of 0.001 as the threshold of significance.

### Cell wall chemistry

Arabidopsis stems from plants just reaching the developmental stage 1 (Altamura et al., 2001) were used to determine lignin and carbohydrate content following a modified Klason method (Porth et al., 2013). Briefly, samples were ground in a Wiley mill to pass a 40-mesh screen, treated with acetone overnight using a Soxhlet apparatus and then dried for 48 h at 50°C. Approximately 100 mg of dried extractive-free tissue was treated with 72% sulphuric acid for 2 h, diluted to ∼3% with 112 ml DI water and autoclaved at 121°C for 60 min. The mixture was filtered through a medium coarseness crucible and the retentate dried at 105°C. The acid-insoluble lignin was determined gravimetrically by weighing the retentate, while the acid-soluble lignin was measured from an aliquot of the filtrate using an UV spectrophotometer at 205 nm. Carbohydrate contents were determined by HPLC analysis of the filtrate. Fucose, glucose, xylose, mannose, galactose, arabinose and rhamnose were analyzed using a Dx-600 anion-exchange HPLC (Dionex, Sunnyvale, CA, USA) equipped with a CarboPac PA1 column (Dionex) at 1 ml min^-1^ and post column detection (100mM NaOH min^-1^) using an electrochemical detector. Sugar concentrations were calculated from standard curves created from external standards.

### Measurements of endogenous cytokinins

Quantification of cytokinin metabolites were performed according to an ultra-high performance liquid chromatography-electrospray tandem mass spectrometry method described by Svačinová et al. (2012). All samples (20 mg FW) were homogenized and extracted in 1 ml of modified Bieleski buffer (60% MeOH, 10% HCOOH and 30% H_2_O) together with a cocktail of stable isotope-labeled internal standards (0.25 pmol of CK bases, ribosides, *N*-glucosides, and 0.5 pmol of CK *O*-glucosides, nucleotides per sample added). The extracts were purified using the Oasis MCX column (30 mg/1 ml, Waters) conditioned with 1 ml each of 100% MeOH and H_2_O, equilibrated sequentially with 1ml of 50% (v/v) nitric acid, 1 ml of H_2_O, and 1 ml of 1M HCOOH. After sample application onto an MCX column, unretained compounds were removed by a wash step using 1 ml of 1M HCOOH and 1 ml 100% MeOH, preconcentrated analytes were eluted by two-step elution using 1 ml of 0.35M NH_4_OH aqueous solution and 2 ml of 0.35M NH_4_OH in 60% (v/v) MeOH solution. The eluates were then evaporated to dryness *in vacuo* and stored at -20°C prior the LC-MS/MS analyses. Cytokinin levels were determined using stable isotope-labelled internal standards as a reference and four independent biological replicates were performed.

### Gene expression analysis

Four stem portions of one cm were collected at the developmental stage when the first silique differentiated on the inflorescence (Altamura et al., 2001). The basal, medial, sub-apical and apical portions of stem correspond respectively to the mature stem above the rosette leaves, the second internode, the third internode (below the last node) and the stem portion just under the apical meristem (Fig. S5A). For each biological replicate, the stem portions from three different plants were pooled and flash frozen immediately in liquid nitrogen. Total RNA was extracted using Trizol reagent (Invitrogen) and genomic DNA was removed by digestion with DNase I (Thermo Scientific) according to the manufacturer’s protocol. One microgram of total RNA was reverse transcribed using the RevertAid First Strand cDNA Synthesis Kit and oligo(dT)18 following the manufacturer’s instructions (Thermo Scientific). Quantitative real-time PCR reaction was performed on a Rotor-Gene Q (Qiagen) using 10 µL FastStart SYBR Green Master (Roche), 2 µL of 2-fold diluted cDNA, and 0.3 µM of primers in a total volume of 20 µL per reaction. The cycling conditions were composed of an initial 10 min denaturation step at 95°C, followed by 45 cycles of 95°C for 10 s, 60°C for 15 s, 72°C for 15 s. A melting curve was run from 65°C to 98°C to ensure the specificity of the products. Data were analyzed with the delta Ct method. Ubiquitin 10 (*UBQ10*) was used as a reference gene for normalization of gene expression levels. qRT-PCR primer sequences are listed in Table S4.

### Statistical analysis

The experiments reported here were repeated at least twice with similar results unless otherwise specified. Experiments were analyzed by ANOVAs followed by Tukey’s honestly significant difference (HSD) *post hoc* test using R software (R Core Team, https://www.R-project.org). For non-normal count data (e.g. number of days) a Poisson mixed model was used to identify differences between genotypes.

## Acknowledgements

We are grateful to Eva Benkova and Phil Jackson for careful reading of the manuscript and their valuable comments, and Hana Martínková and Ivan Petřík for their help with phytohormone analyses. Plant Sciences Core Facility of CEITEC Masaryk University is gratefully acknowledged for plants cultivation.

## Data availability

All relevant data can be found within the article and its supplementary information.

## References

1. Altamura, M. M., Possenti, M., Matteucci, A., Baima, S., Ruberti, I. and Morelli, G. (2001). Development of the vascular system in the inflorescence stem of Arabidopsis. New Phytologist 151, 381–389.

2. Argyros, R. D., Mathews, D. E., Chiang, Y. H., Palmer, C. M., Thibault, D. M., Etheridge, N., Argyros, D. A., Mason, M. G., Kieber, J. J. and Schaller, G. E. (2008). Type B response regulators of Arabidopsis play key roles in cytokinin signaling and plant development. Plant Cell 20, 2102–2116.

3. Baima, S., Possenti, M., Matteucci, A., Wisman, E., Altamura, M. M., Ruberti, I. and Morelli, G. (2001). The arabidopsis ATHB-8 HD-zip protein acts as a differentiation-promoting transcription factor of the vascular meristems. Plant Physiol 126, 643–655.

4. Braybrook, S. A. and Jonsson, H. (2016). Shifting foundations: the mechanical cell wall and development. Curr Opin Plant Biol 29, 115–120.

5. Campbell, L., Etchells, J. P., Cooper, M., Kumar, M. and Turner, S. R. (2018). An essential role for abscisic acid in the regulation of xylem fibre differentiation. Development 145.

6. Dello Ioio, R., Linhares, F. S. and Sabatini, S. (2008). Emerging role of cytokinin as a regulator of cellular differentiation. Curr Opin Plant Biol 11, 23–27.

7. Didi, V., Jackson, P. and Hejatko, J. (2015). Hormonal regulation of secondary cell wall formation. Journal of Experimental Botany 66, 5015–5027.

8. Eden, E., Navon, R., Steinfeld, I., Lipson, D. and Yakhini, Z. (2009). GOrilla: a tool for discovery and visualization of enriched GO terms in ranked gene lists. BMC Bioinformatics 10, 48.

9. Esau, K. (1977). Anatomy of seed plants. New York: Wiley.

10. Gajdosova, S., Spichal, L., Kaminek, M., Hoyerova, K., Novak, O., Dobrev, P. I., Galuszka, P., Klima, P., Gaudinova, A., Zizkova, E., et al. (2011). Distribution, biological activities, metabolism, and the conceivable function of cis-zeatin-type cytokinins in plants. J Exp Bot 62, 2827–2840.

11. Gordon, S. P., Chickarmane, V. S., Ohno, C. and Meyerowitz, E. M. (2009). Multiple feedback loops through cytokinin signaling control stem cell number within the Arabidopsis shoot meristem. Proc Natl Acad Sci U S A 106, 16529–16534.

12. Greb, T. and Lohmann, J. U. (2016). Plant Stem Cells. Curr Biol 26, R816–821.

13. Hejatko, J., Ryu, H., Kim, G. T., Dobesova, R., Choi, S., Choi, S. M., Soucek, P., Horak, J., Pekarova, B., Palme, K., et al. (2009). The Histidine Kinases CYTOKININ-INDEPENDENT1 and ARABIDOPSIS HISTIDINE KINASE2 and 3 Regulate Vascular Tissue Development in Arabidopsis Shoots. Plant Cell 21, 2008–2021.

14. Higuchi, M., Pischke, M. S., Mahonen, A. P., Miyawaki, K., Hashimoto, Y., Seki, M., Kobayashi, M., Shinozaki, K., Kato, T., Tabata, S., et al. (2004). In planta functions of the Arabidopsis cytokinin receptor family. Proc Natl Acad Sci U S A 101, 8821–8826.

15. Hosek, P., Hoyerova, K., Kiran, N. S., Dobrev, P. I., Zahajska, L., Filepova, R., Motyka, V., Muller, K. and Kaminek, M. (2019). Distinct metabolism of N-glucosides of isopentenyladenine and trans-zeatin determines cytokinin metabolic spectrum in Arabidopsis. New Phytol.

16. Hussey, S. G., Mizrachi, E., Creux, N. M. and Myburg, A. A. (2013). Navigating the transcriptional roadmap regulating plant secondary cell wall deposition. Front Plant Sci 4, 325.

17. Chebli, Y. and Geitmann, A. (2017). Cellular growth in plants requires regulation of cell wall biochemistry. Curr Opin Cell Biol 44, 28–35.

18. Jung, K. W., Oh, S. I., Kim, Y. Y., Yoo, K. S., Cui, M. H. and Shin, J. S. (2008). Arabidopsis histidine-containing phosphotransfer factor 4 (AHP4) negatively regulates secondary wall thickening of the anther endothecium during flowering. Mol Cells 25, 294–300.

19. Jupa, R., Didi, V., Hejatko, J. and Gloser, V. (2015). An improved method for the visualization of conductive vessels in Arabidopsis thaliana inflorescence stems. Front Plant Sci 6, 211.

20. Kieber, J. J. and Schaller, G. E. (2018). Cytokinin signaling in plant development. Development 145.

21. Kim, D., Pertea, G., Trapnell, C., Pimentel, H., Kelley, R. and Salzberg, S. L. (2013). TopHat2: accurate alignment of transcriptomes in the presence of insertions, deletions and gene fusions. Genome Biol 14, R36.

22. Kubo, M., Udagawa, M., Nishikubo, N., Horiguchi, G., Yamaguchi, M., Ito, J., Mimura, T., Fukuda, H. and Demura, T. (2005). Transcription switches for protoxylem and metaxylem vessel formation. Gene Dev 19, 1855–1860.

23. Liu, C., Yu, H., Rao, X., Li, L. and Dixon, R. A. (2021). Abscisic acid regulates secondary cell-wall formation and lignin deposition in Arabidopsis thaliana through phosphorylation of NST1. Proc Natl Acad Sci U S A 118.

24. Martin, M. (2011). Cutadapt removes adapter sequences from high-throughput sequencing reads. EMBnet.journal 17, 10–12.

25. Matsumoto-Kitano, M., Kusumoto, T., Tarkowski, P., Kinoshita-Tsujimura, K., Vaclavikova, K., Miyawaki, K. and Kakimoto, T. (2008). Cytokinins are central regulators of cambial activity. Proc Natl Acad Sci U S A 105, 20027–20031.

26. Mitsuda, N., Iwase, A., Yamamoto, H., Yoshida, M., Seki, M., Shinozaki, K. and Ohme-Takagi, M. (2007). NAC transcription factors, NST1 and NST3, are key regulators of the formation of secondary walls in woody tissues of Arabidopsis. Plant Cell 19, 270-280.

27. Mitsuda, N., Seki, M., Shinozaki, K. and Ohme-Takagi, M. (2005). The NAC transcription factors NST1 and NST2 of Arabidopsis regulate secondary wall thickenings and are required for anther dehiscence. The Plant cell 17, 2993–3006.

28. Miyawaki, K., Tarkowski, P., Matsumoto-Kitano, M., Kato, T., Sato, S., Tarkowska, D., Tabata, S., Sandberg, G. and Kakimoto, T. (2006). Roles of Arabidopsis ATP/ADP isopentenyltransferases and tRNA isopentenyltransferases in cytokinin biosynthesis. Proc Natl Acad Sci U S A 103, 16598–16603.

29. Nieminen, K., Immanen, J., Laxell, M., Kauppinen, L., Tarkowski, P., Dolezal, K., Tahtiharju, S., Elo, A., Decourteix, M., Ljung, K., et al. (2008). Cytokinin signaling regulates cambial development in poplar. Proc Natl Acad Sci U S A 105, 20032–20037.

30. Nilsson, J., Karlberg, A., Antti, H., Lopez-Vernaza, M., Mellerowicz, E., Perrot-Rechenmann, C., Sandberg, G. and Bhalerao, R. P. (2008). Dissecting the molecular basis of the regulation of wood formation by auxin in hybrid aspen. Plant Cell 20, 843–855.

31. Nishimura, C., Ohashi, Y., Sato, S., Kato, T., Tabata, S. and Ueguchi, C. (2004). Histidine kinase homologs that act as cytokinin receptors possess overlapping functions in the regulation of shoot and root growth in Arabidopsis. Plant Cell 16, 1365–1377.

32. Pernisova, M., Klima, P., Horak, J., Valkova, M., Malbeck, J., Soucek, P., Reichman, P., Hoyerova, K., Dubova, J., Friml, J., et al. (2009). Cytokinins modulate auxin-induced organogenesis in plants via regulation of the auxin efflux. Proc Natl Acad Sci U S A 106, 3609–3614.

33. Porth, I., Klapste, J., Skyba, O., Lai, B. S., Geraldes, A., Muchero, W., Tuskan, G. A., Douglas, C. J., El-Kassaby, Y. A. and Mansfield, S. D. (2013). Populus trichocarpa cell wall chemistry and ultrastructure trait variation, genetic control and genetic correlations. New Phytol 197, 777–790.

34. Ramachandran, P., Augstein, F., Mazumdar, S., Nguyen, T. V., Minina, E. A., Melnyk, C. W. and Carlsbecker, A. (2021). Abscisic acid signaling activates distinct VND transcription factors to promote xylem differentiation in Arabidopsis. Curr Biol 31, 3153–3161 e3155.

35. Riefler, M., Novak, O., Strnad, M. and Schmulling, T. (2006). Arabidopsis cytokinin receptor mutants reveal functions in shoot growth, leaf senescence, seed size, germination, root development, and cytokinin metabolism. Plant Cell 18, 40–54.

36. Ruzicka, K., Ursache, R., Hejatko, J. and Helariutta, Y. (2015). Xylem development - from the cradle to the grave. New Phytol 207, 519–535.

37. Sassi, M. and Traas, J. (2015). When biochemistry meets mechanics: a systems view of growth control in plants. Curr Opin Plant Biol 28, 137–143.

38. Schafer, M., Brutting, C., Meza-Canales, I. D., Grosskinsky, D. K., Vankova, R., Baldwin, I. T. and Meldau, S. (2015). The role of cis-zeatin-type cytokinins in plant growth regulation and mediating responses to environmental interactions. J Exp Bot 66, 4873–4884.

39. Schuetz, M., Smith, R. and Ellis, B. (2013). Xylem tissue specification, patterning, and differentiation mechanisms. Journal of Experimental Botany 64, 11–31.

40. Skalak, J., Nicolas, K. L., Vankova, R. and Hejatko, J. (2021). Signal Integration in Plant Abiotic Stress Responses via Multistep Phosphorelay Signaling. Front Plant Sci 12, 644823.

41. Svacinova, J., Novak, O., Plackova, L., Lenobel, R., Holik, J., Strnad, M. and Dolezal, K. (2012). A new approach for cytokinin isolation from Arabidopsis tissues using miniaturized purification: pipette tip solid-phase extraction. Plant Methods 8, 17.

42. Taylor-Teeples, M., Lin, L., de Lucas, M., Turco, G., Toal, T. W., Gaudinier, A., Young, N. F., Trabucco, G. M., Veling, M. T., Lamothe, R., et al. (2015). An Arabidopsis gene regulatory network for secondary cell wall synthesis. Nature 517, 571-575.

43. Trapnell, C., Hendrickson, D. G., Sauvageau, M., Goff, L., Rinn, J. L. and Pachter, L. (2013). Differential analysis of gene regulation at transcript resolution with RNA-seq. Nat Biotechnol 31, 46–53.

44. Trinh, D. C., Alonso-Serra, J., Asaoka, M., Colin, L., Cortes, M., Malivert, A., Takatani, S., Zhao, F., Traas, J., Trehin, C., et al. (2021). How Mechanical Forces Shape Plant Organs. Curr Biol 31, R143–R159.

45. Tyree, M. T., Davis, S. D. and Cochard, H. (1994). Biophysical Perspectives of Xylem Evolution - Is There a Tradeoff of Hydraulic Efficiency for Vulnerability to Dysfunction. Iawa J 15, 335–360.

46. Ye, L., Wang, X., Lyu, M., Siligato, R., Eswaran, G., Vainio, L., Blomster, T., Zhang, J. and Mahonen, A. P. (2021). Cytokinins initiate secondary growth in the Arabidopsis root through a set of LBD genes. Curr Biol 31, 3365–3373 e3367.

47. Zhong, R., Richardson, E. A. and Ye, Z. H. (2007). Two NAC domain transcription factors, SND1 and NST1, function redundantly in regulation of secondary wall synthesis in fibers of Arabidopsis. Planta 225, 1603–1611.

48. Zhong, R. and Ye, Z. H. (2015). Secondary cell walls: biosynthesis, patterned deposition and transcriptional regulation. Plant Cell Physiol 56, 195–214.

49. Zhou, J. L., Zhong, R. Q. and Ye, Z. H. (2014). Arabidopsis NAC Domain Proteins, VND1 to VND5, Are Transcriptional Regulators of Secondary Wall Biosynthesis in Vessels. Plos One 9, 13.

50. Zubo, Y. O. and Schaller, G. E. (2020). Role of the Cytokinin-Activated Type-B Response Regulators in Hormone Crosstalk. Plants (Basel) 9.

